# Norepinephrine Signals Through Astrocytes To Modulate Synapses

**DOI:** 10.1101/2024.05.21.595135

**Authors:** Katheryn B Lefton, Yifan Wu, Allen Yen, Takao Okuda, Yufen Zhang, Yanchao Dai, Sarah Walsh, Rachel Manno, Joseph D Dougherty, Vijay K Samineni, Paul C Simpson, Thomas Papouin

## Abstract

Locus coeruleus (LC)-derived norepinephrine (NE) drives network and behavioral adaptations to environmental saliencies by reconfiguring circuit connectivity, but the underlying synapse-level mechanisms are elusive. Here, we show that NE remodeling of synaptic function is independent from its binding on neuronal receptors. Instead, astrocytic adrenergic receptors and Ca^2+^ dynamics fully gate the effect of NE on synapses as the astrocyte-specific deletion of adrenergic receptors and three independent astrocyte-silencing approaches all render synapses insensitive to NE. Additionally, we find that NE suppression of synaptic strength results from an ATP-derived and adenosine A1 receptor-mediated control of presynaptic efficacy. An accompanying study from Chen et al. reveals the existence of an analogous pathway in the larval zebrafish and highlights its importance to behavioral state transitions. Together, these findings fuel a new model wherein astrocytes are a core component of neuromodulatory systems and the circuit effector through which norepinephrine produces network and behavioral adaptations, challenging an 80-year-old status quo.

## Introduction

Neuromodulators exert global control over brain function and behavior by reshaping brain activity and functional connectivity. A canonical example, norepinephrine (NE), is produced by locus coeruleus (LC) neurons that extend NE-releasing projections throughout the brain (*1, 2*). The activity of the LC-NE system is broadly associated with network and behavioral adaptations in response to environmental contingencies, making it instrumental to brain state transitions, processing of sensory saliencies, as well as cognitive functions including goal-directed decision-making, learning, and cognitive flexibility (*3, 4*). The significance of LC-derived NE signaling to behavior relates to its ability to functionally reorganize neural circuits by altering the strength of synaptic connections within networks, with original evidence dating back eight decades (*5–8*). This is central to mesoscale theories that bridge the behavioral and cellular effects of the LC-NE system, such as the ‘adaptive gain’ (*9*), ‘network reset’ (*10*), and ‘global model failure’ theories (*11*). But, in contrast with their broad applicability, these models conceal a surprisingly poor understanding of the effects of NE on synapses.

A broadly accepted view is that NE remodels synaptic networks by acting on cognate receptors on neurons (*1, 3*), but this is now at odds with a growing appreciation for the multicellular nature of the brain and evidence that NE also signals onto non-neuronal cells such as astrocytes (*12, 13*). Astrocytes are ubiquitous circuit components of the central nervous systems (CNS) of vertebrates, and other bilateria. They each form an extensive meshwork of ultrafine processes known to infiltrate and control the microenvironment of 10^5^ synapses, and other functional units, and they are increasingly recognized as state-dependent orchestrators of neural circuit function (*14–17*). Astrocytes express various noradrenergic receptors (*18, 19*) and respond to NE occurrence with cell-wide elevations of intracellular free calcium (Ca^2+^), a phenomenon observed across phylogenetically distant species (*20–23*). Ca^2+^-dependent intracellular cascades, in turn, mobilize various forms of astrocyte activities and outputs that are potent regulators of synaptic and network function (*20, 22, 24*). Yet, whether astrocytes actively contribute to the circuit effects of neuromodulators such as NE remains an open question.

## Results

### LC-derived NE reduces synaptic efficacy

The ability of NE to reshape synaptic connectivity has been documented in diverse species, preparations, and regions of the CNS (*3, 8, 25–35*). In order to interrogate underlying mechanisms in a tractable system, we performed extracellular recordings of AMPAR-mediated field excitatory postsynaptic potentials (fEPSPs) in the *stratum radiatum* of acute hippocampal slices taken from adult mice (Fig.1A, Methods and SupFig.1A). In accord with decades of work, we found that the bath application of NE produced a marked and concentration-dependent decrease in synaptic strength (fEPSP slope, Fig.1B,C, SupFig.1B. All statistical analyses are reported in Table S1). This rapid remodeling was distinct from the NMDAR-mediated, activity-dependent long-term depression caused by NE (*34, 36*) because it persisted in the presence of the NMDAR blocker D-AP5 and was still observed when synaptic stimulations were paused at the onset of NE application (SupFig.1C-E). Importantly, the magnitude of NE-induced inhibition did not depend on initial synaptic properties (SupFig.1F) which, together with the presence of GABA_A_-R blocker in the bath, ruled out the possibility of an inhibitory feedback mechanism. Changes in presynaptic properties were simultaneously assessed with a classic paradigm consisting of pairs of stimulations, delivered 200ms apart. This yielded a paired-pulse facilitation index (PPF, Methods), which varies inversely with the probability of neurotransmitter release at presynaptic terminals. Concomitant to its effect on synaptic strength, NE elicited an increase in PPF, indicative of a reduction in release probability (Fig.1B,C). The increase in PPF and decline in fEPSP slope closely coincided in their temporal profiles (Fig.1B), and their magnitude strongly correlated (Fig.1D), pointing at a primarily presynaptic site of action of NE. This was directly confirmed using minimal-stimulation experiments in whole-cell patch-clamp recordings, which allow monitoring unitary excitatory postsynaptic currents (EPSCs) occurring at single synapses (Fig.1A,E, Methods) (*37*). Under these conditions, an increased rate of presynaptic failure became apparent within minutes of NE application (i.e., decreased (pre)synaptic efficacy, Fig.1E-G), with no change in the amplitude of successful EPSCs (i.e., (post)synaptic potency), demonstrating that NE inhibits synapse strength (total EPSC amplitude) via a presynaptic mechanism.

**Fig. 1:**
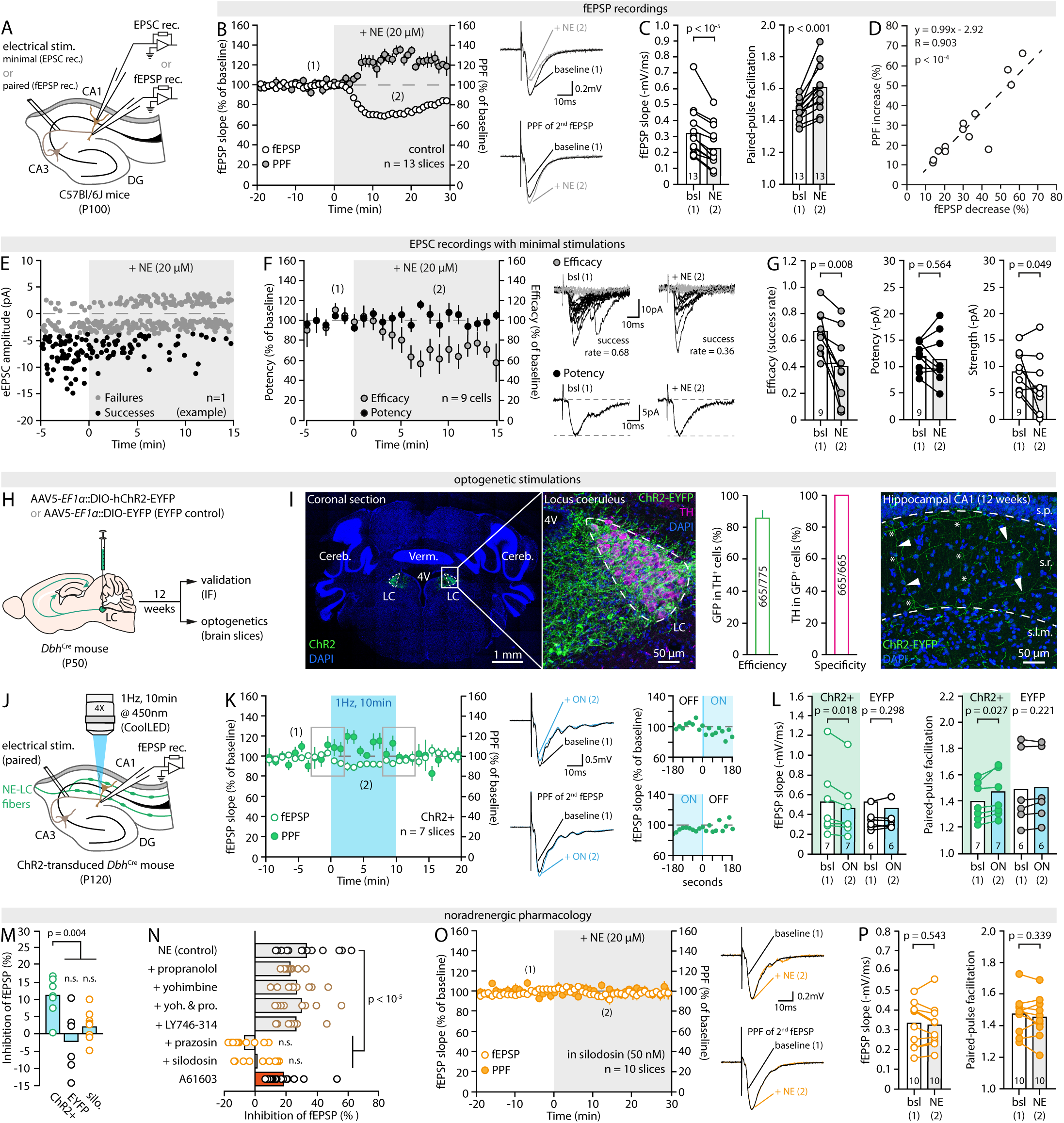
NE and LC-NE activity inhibit presynaptic efficacy. (**A**) Schematic of the recording conditions. (**B**) *Left*, Time-course of the effect of 20µM NE (applied at t=0, greyed area) on fEPSP slope and PPF. Each circle is the average of three data points per minute. (1) and (2) indicate the approximate epochs at which sample traces were obtained and quantifications performed for the baseline and NE conditions. *Right*, Representative traces showing the effect of NE on the slope of the first fEPSP (top) and on the PPF of the second fEPSP (bottom) from a same recording. Stimulation artefacts were cropped for clarity. (**C**) Pair-wise quantification of the effect of NE on fEPSP slope and PPF for the experiments shown in (B). (**D**) Correlation between NE-induced fEPSP decrease and PPF increase in experiments shown in (B) and (C). (**E**) Representative recording of EPSCs amplitude in response to minimal stimulations, showing failures (grey) and successes (black). (**F**) *Left*, Averaged time course (per minute) of minimal-stimulation experiments showing the effect of 20µM NE on synaptic efficacy (grey) and potency (black). *Right*, Representative traces illustrating the effect of NE on efficacy (grey traces, failures; black traces, successes) and potency (average of successes) over 1min epochs. (**G**) Quantification and pair-wise comparison of the effect of NE on synaptic efficacy, potency and strength for individual experiments shown in (F). (**H**) Schematic illustration of the experimental procedure for expressing ChR2 in LC-NE fibers. (**I**) IHC images of ChR2-EYFP expression in the LC (*left*) and hippocampal CA1 (*right*) at 12 weeks, and quantification of efficiency and specificity of ChR2-EYFP expression in the LC (*center*). Tyrosine hydroxylase (TH) is used as a marker of NE-producing neurons (n= 22 sections, 4 animals). Cereb.: Cerebellum; 4V: fourth ventricle; Verm.: Vermis; s.p.: stratum pyramidale, s.r.: stratum radiatum; s.l.m.: stratum lacunasorum moleculare. Arrowheads indicate ChR2-EYFP-expressing LC-NE projections with major bifurcation points (asterisks). (**J**) Schematic of the recording and optogenetic stimulation conditions. (**K**) Time-course of the effect of the optical stimulation of LC-NE fibers on fEPSPs and PPF, and representative traces. Insets show the detailed time-course (20s bins) at the onset and offset of light (s.e.m. omitted for clarity). (**L**) Pair-wise quantification of the effect of light (1Hz, 10min) on fEPSP slope and PPF for the experiments shown in (K) and in EYFP-control slices. (**M**) Plot summarizing the effect of light in ChR2-positive slices, EYFP-control slices, and ChR2-positive slices in the presence of silodosin (50nM, silo.). (**N**) Plot summarizing the effect of NE on synaptic strength in the presence of blockers of ɑ2-AR (yohimbine, 500nM), β-AR (propranolol, 1µM), ɑ1B-AR (LY746-314, 1µM), ɑ1-AR (prazosin, 1µM), ɑ1A-AR (silodosin, 50nM) or the ɑ1A-AR agonist A61603 (70nM). (**O,P**) Time-course of the effect of 20µM NE on fEPSPs and PPF in the presence of silodosin (50nM), representative traces, and pair-wise quantification. Data are shown as mean ± s.e.m.

We next set out to verify that these observations were not limited to the bath application of exogenous NE, using optogenetics. To drive the specific expression of channel-rhodopsin (ChR2) in LC-NE neurons, we micro-injected AAV5-*EF1α*::DIO-hChR2(H134R)-EYFP into the LC of *Dbh*^Cre^ knock-in mice expressing Cre recombinase from the endogenous dopamine beta hydroxylase locus (*38*), which encodes an oxidoreductase that catalyzes the formation of NE from dopamine (*39*) (Fig.1H,I). Abundant expression of EYFP in LC-NE fibers that densely innervate the hippocampal CA1 became evident 10 weeks later and plateaued at 12 weeks (*38, 40*) (Fig.1I and SupFig.1G). Acute hippocampal slices were obtained 12 weeks post-injection (Methods), and fEPSPs and PPF were recorded as above. LC-NE fibers fire tonically at 1-3Hz in vivo, with bouts of 10Hz phasic activity, and are often experimentally stimulated with paradigms as strong as 25Hz (*38, 41*). We found that an optical stimulation as weak as 1 Hz (450 nm, 15mW, 10min, Fig.1J) caused a modest but reproducible and sustained decline in fEPSP slope accompanied by an equivalent increase in PPF (Fig.1K,L), neither of which were observed in EYFP control slices taken from AAV5-*EF1α*::DIO-EYFP injected *Dbh*^Cre^ mice (Fig.1L,M and SupFig.1H, and Table S1). These effects appeared within 20sec of light onset, remained constant for the duration of the stimulation, and subsided within 2min of light cessation (Fig.1K). Taken together, these results support the notion that exogenous and endogenous NE inhibits transmission at excitatory synapses by hindering (pre)synaptic efficacy.

### LC-derived NE alters synapses via ɑ1-ARs

NE binds three classes of G-protein coupled receptors (GPCRs): alpha1 adrenergic receptors (ɑ1-AR), alpha2 AR (ɑ2-AR) and beta AR (β-ARs). Consistent with prior work, we found that the ɑ1-AR antagonist prazosin blocked the effect of NE both at commonly used doses (10µM, SupFig.1K) and at concentrations no greater than ∼100 times its pKi for ɑ1-ARs (1µM, Fig.1N, SupFig.1L) (*42*). Conversely, ɑ2-AR and β-AR antagonists, yohimbine and propranolol, respectively, alone or in combination, failed to block the effect of NE even at higher doses (Fig.1N and SupFig.1M-P). Additionally, we found that NE applications (Fig.1N-P) and optogenetic stimulations of LC-NE fibers (Fig.1M and SupFig.1I,J) were ineffective in the presence of the ɑ1A-AR subtype-specific antagonist silodosin (50nM), but unaffected by LY746-314, an ɑ1B-AR specific antagonist (1µM, Fig.1N and SupFig.1Q). Furthermore, the ɑ1A-AR agonist A61603 mimicked the effect of NE (70nM, Fig.1N and SupFig.1R,S). Hence, in our conditions, the action of NE depends on the ɑ1A-AR subtype alone.

While in line with existing literature (*33, 34*), that NE suppresses presynaptic release probability via ɑ1A-ARs is surprising in two ways. First, ɑ1A-ARs are G_q_-coupled GPCRs, hence their activation should enhance Ca^2+^ influx in synaptic boutons and facilitate synaptic transmission (*43*). Second, the IC_50_ of the observed effect of NE on synapses (3.5µM, SupFig.1B) is 12 times greater than its binding affinity on the ɑ1A-AR (∼0.28µM (*44*)). In total, the presynaptic localization, affinity, and mode of action of ɑ1A-AR are thus hard to reconcile with our observations, hinting at the possibility that NE alters synaptic function through a more intricate mechanism than commonly assumed.

### Astrocyte Ca^2+^ dynamics gate NE effectiveness

A recent paradigm shift in neuroscience has been the realization that synaptic connectivity is the result of a subtle multicellular interplay between fast neuronal activity and instructive or permissive signals from non-neuronal cells, including astrocytes (*45*). To capture the effect of NE on astrocytes, we transduced C57BL/6J adult mice (P70) with an AAV5-*gfaABC1D*::lck-GCaMP6f in the CA1 region of the hippocampus (SupFig.2A), yielding robust expression of the Ca^2+^ indicator lck-GCaMP6f in GFAP-positive cells (astrocytes) of the *stratum radiatum* (SupFig.2B), and performed two-photon laser scanning microscopy imaging (2-PLSM) in acute slices 20-30 days later (SupFig.2C,D, and Methods). Consistent with previous reports (*18, 20*), NE applications elicited large, cell-wide and dose-dependent Ca^2+^ responses in nearly all astrocytes in the field of view (SupFig.2E). Strikingly, the ɑ1A-AR specific antagonist silodosin (50nM), which obliterated the effect of NE on synapses (Fig.1N-P), markedly hampered this response (SupFig.2F). In addition, silodosin, in itself, reduced the amplitude and frequency of astrocyte spontaneous Ca^2+^ transients (SupFig.2G,H), while it increased fEPSP slope and reduced PPF (SupFig.1T), indicating that a tonic ɑ1A-AR-mediated activation of astrocytes coincided with tonic ɑ1A-AR-mediated inhibition of synapses. Combined, these observations hinted at a causal link between astrocyte Ca^2+^ dynamics and synaptic changes caused by NE.

To test this idea, we employed iβARK, a tool that prevents Ca^2+^ elevations secondary to the activation of G_q_-GPCRs in astrocytes (*46*). The transduction of AAV5-*gfaABC1D*::iβARK-mCherry in the CA1 of wild-type mice (Fig.2A) yielded a robust, cell-specific expression in 92% of hippocampal astrocytes (Fig.2B). Basal evoked fEPSPs amplitude and PPF were normal in slices obtained from iβARK-transduced animals (iβARK-slices, Fig.2C) and spontaneous Ca^2+^ dynamics were virtually unchanged in iβARK-expressing astrocytes (Fig.2D,E). NE-evoked Ca^2+^ responses, however, were blunted by 75% in iβARK-expressing astrocytes relative to AAV5-*gfaABC1D*::mCherry-transduced astrocytes (RFP-control, Fig.2F,G), consistent with their reliance on G_q_-coupled ɑ1A-ARs (SupFig.2). Remarkably, NE yielded the expected effect on synapses in RFP-control slices (Fig.2H) but only elicited a marginal and statistically insignificant decline of fEPSP slope in iβARK-slices, with no noticeable effect on the PPF (Fig.2I,J). Interestingly, while greatly reduced compared to the RFP-control (Fig.2K), the impact of NE remained evident in some iβARK-slices. It averaged 4.8% across experiments, equivalent to 21% of the inhibition achieved in RFP-controls (Fig.2K), which matches the proportion of NE-induced Ca^2+^ response that persisted in iβARK-astrocytes (∼20% of RFP-controls, Fig.2G). It is still unknown whether Ca^2+^ input-output coding in astrocytes follows a threshold-based all-or-nothing logic, correlates linearly with the magnitude of Ca^2+^ events, or follows other nonlinear rules, muddying the rigorous interpretation of these data. Remarkably, however, we obtained identical results with other proven methods of astrocyte Ca^2+^-silencing, such as CalEx (calcium extrusion, Fig.2L and SupFig.3) (*47*) and thapsigargin (Fig.2L and SupFig.4) (*48*). Because these approaches are mechanistically distinct, we interpret the apparent residual effect on synaptic strength in all three conditions as the consequence of the incomplete blockade of NE-induced astrocyte Ca^2+^ elevations achieved by each of them, and we conclude that silencing astrocytes occludes NE neuromodulation of synapses. An alternative interpretation, however, is that astrocytes contribute the majority (70-80%), but not all the inhibitory effect of NE, with the rest (20-30%) being attributable to a direct action on neurons.

**Fig. 2:**
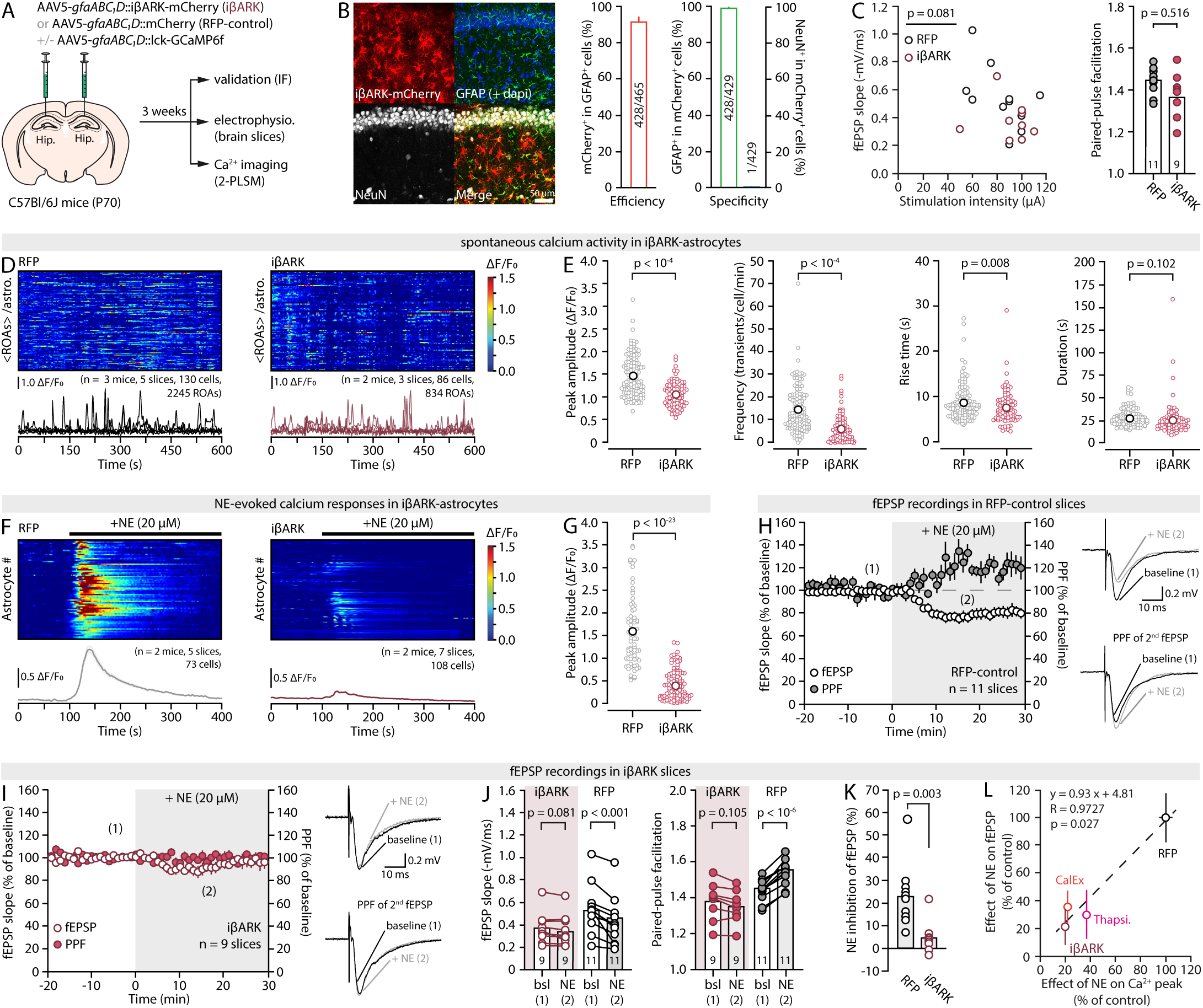
Astrocyte Ca^2+^ dynamics gate the effect of NE on synapses. (**A**) Approach for astrocyte Ca^2+^ silencing with iβARK. (**B**) Representative IHC images of iβARK-mCherry expression in the hippocampal CA1, along with quantification of efficiency and specificity (n=5 slices). (**C**) Plot of the stimulation intensity/fEPSP slope relationship (*left*, unpaired Student’s *t*-test on slope/stim ratio) and summary bar graphs of PPF values (*right*) in RFP-control and iβARK slices at baseline. (**D**) Kymograph (each row shows the average fluorescence across ROAs of a single astrocyte) and 5 representative ΔF/F_0_ traces (from individual ROAs) of spontaneous Ca^2+^ transients in RFP-control and iβARK slices. Horizontal time axis applies to the kymograph and representative traces. (**E**) Pots of the peak amplitude, frequency, and kinetics of spontaneous Ca^2+^ transients in RFP-control and iβARK slices. Each data point shows the average fluorescence across ROAs for an individual astrocyte. (**F**) Kymographs (each row shows the whole-cell fluorescence of a single astrocyte) and average ΔF/F_0_ traces (± s.e.m.) across all astrocytes in response to 20µM NE application in RFP-control and iβARK slices. (**G**) Plot of the peak ΔF/F_0_ response in RFP-control and iβARK conditions for experiments shown in (F). (**H,I**) Time-courses of the effect of 20µM NE on fEPSP slope and PPF, and representative traces, in RFP-control and iβARK slices. (**J**) Pair-wise quantifications of the effect of NE on fEPSP slope and PPF for the experiments shown in (H) and (I). (**K**) Plots summarizing the effect of NE on fEPSP slope in RFP-control and iβARK slices. (**L**) Correlation between the effect of 20µM NE on astrocyte peak Ca^2+^ responses and fEPSP slope across three methods of astrocyte silencing and RFP-controls (see SupFig.3, SupFig.4 and Table S1). Data are shown as mean ± s.e.m.

### Neuronal ɑ1-ARs are not required for NE neuromodulation

To parse out how direct signaling of NE onto neurons and indirect signaling through astrocytes contribute to the functional remodeling of synapses, we sought to delete ɑ1A-AR from pre-synaptic, post-synaptic and inhibitory neurons by micro-injecting AAV5-*hSyn*::Cre-GFP in the hippocampus of *Adra1a^fl/fl^* mice (Fig.3A). Immuno-fluorescent examinations confirmed the strong expression of the GFP reporter in 88% and 89% of CA3 and CA1 NeuN-positive cells (neurons), respectively, with extremely high neuronal specificity (Fig.3B). Additionally, RT-PCR showed a 46% decrease of unmodified *Adra1a* gDNA in GFP-positive cells compared to GFP-negative in Cre-transduced *Adra1a^fl/fl^*hippocampi (Fig.3A,C and Methods), together confirming the effective ablation of ɑ1A-ARs from most CA3 (presynaptic) and CA1 (postsynaptic) neurons in Cre-injected *Adra1a^fl/fl^*mice (neuron-specific *Adra1a* knockout, N-Adra1a^KO^). The inhibitory action of NE on synapses was next assessed in N-Adra1a^KO^ slices, using AAV5-*hSyn*::Cre-GFP injected *Adra1a^+/+^* animals as controls (N-Adra1a^Cre-GFP^). Strikingly, NE yielded a sharp decline in fEPSPs and increase in PPF in N-Adra1a^Cre-GFP^ as well as N-Adra1a^KO^ slices (Fig.3D,E and SupFig.5B). Both conditions were statistically indistinguishable from one-another (Fig.3F, Table S1). Additionally, ɑ1A-AR deletion, on its own, did not alter synaptic strength or pre-synaptic release probability (SupFig.5A). In total, the deletion of ɑ1A-AR from neurons was thus without effect on basal synaptic properties or NE-induced remodeling of synaptic function.

**Fig. 3:**
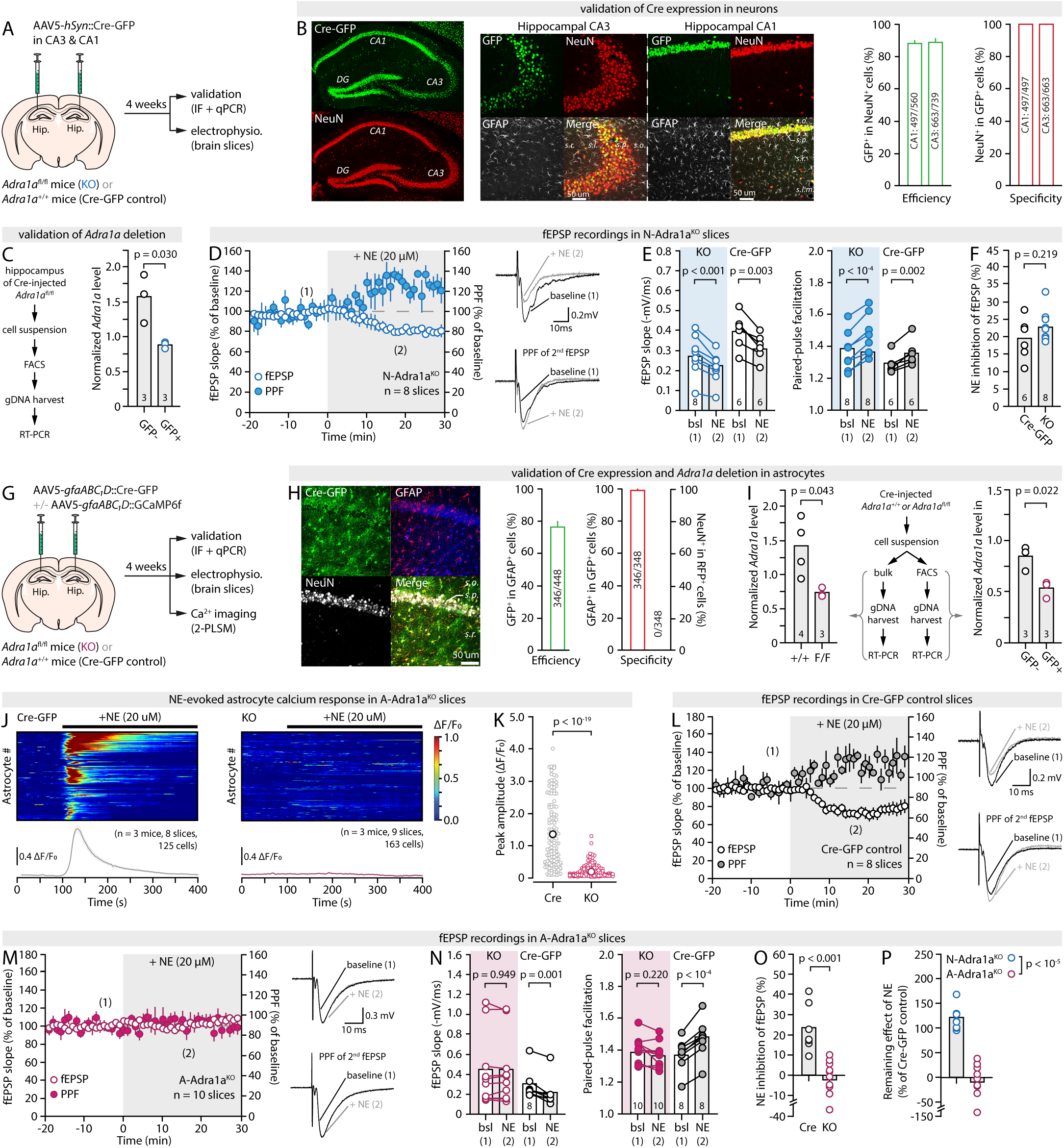
Astrocytic, but not neuronal α1A-ARs are required for NE to affect synapses. (**A**) Approach for the neuronal deletion of *Adra1a*. (**B**) Representative IHC images of Cre-GFP expression in hippocampal CA1 and CA3 neurons, along with quantification of efficiency and specificity (n= 4 sections, 2 animals). (**C**) Quantification of *Adra1a* gDNA levels, normalized to *Atcb* (β-actin), in GFP+ and GFP-cells sorted from N-Adra1a^KO^ animal hippocampi (n = 3). (**D**) Time-course of the effect of 20µM NE on fEPSPs and PPF in N-Adra1a^KO^ slices and representative traces. (**E-F**) Pair-wise quantification of the effect of NE on fEPSP slope and PPF in N-Adra1a^KO^ and N-Adra1a^Cre-GFP^ control slices, and summary plot of the effect of NE on fEPSPs in both conditions. (**G**) Approach for the astrocytic deletion of *Adra1a*. (**H**) Representative IHC images of Cre-GFP expression in hippocampal CA1 astrocytes, along with quantification of efficiency and specificity (n = 6 sections, 2 animals). (**I**) *Left*, quantification of bulk *Adra1a* gDNA levels, normalized to *Actb* (β-actin), in hippocampi from A-Adra1a^Cre-GFP^ controls (n = 4) and A-Adra1a^KO^ (n = 3). *Right*, quantification of *Adra1a* gDNA levels, normalized to *Actb* (β-actin), in GFP+ and GFP-cells sorted from A-Adra1a^KO^ hippocampi (n = 3). (**J,K**) Kymographs (each row shows the whole-cell fluorescence of a single astrocyte), average ΔF/F_0_ traces (± s.e.m.) across all astrocytes, and quantification of the peak Ca^2+^ signal in response to 20µM NE application in A-Adra1a^Cre-GFP^ and A-Adra1a^KO^ slices. (**L-O**) Time-courses of the effect of NE on fEPSPs and PPF, and representative traces, in A-Adra1a^control^ (L) and A-Adra1a^KO^ slices (M), pair-wise quantification of the effect of NE on fEPSP slope and PPF (N), and summary plot of the effect of NE in both conditions (O). **(P)** Summary plot of the inhibitory effect of NE on fEPSPs, relative to controls, in A-Adra1a^KO^ and N-Adra1a^KO^ slices.

### Astrocytic ɑ1-AR deletion renders NE inoperant

The above results imply that astrocytes mediate the entirety of the modulatory action of NE on synapses. To test this, ɑ1A-ARs were deleted from astrocytes specifically, by transducing AAV5-*gfaABC1D*::Cre-GFP (*12*) in the CA1 of *Adra1a^fl/fl^* mice (Fig.3G). Robust expression of GFP was apparent in 76% of *stratum radiatum* astrocytes with 99% specificity 4 weeks later (Fig.3H), accompanied by a 37% reduction of unmodified *Adra1a* gDNA in fluorescence-sorted GFP-positive cells relative to GFP-negative (Fig.3I), and a 51% reduction in bulk *Adra1a* gDNA in hippocampi from AAV5-*gfaABC1D*::Cre-GFP-transduced *Adra1a^fl/fl^* mice (astrocyte-specific *Adra1a* knockouts, A-Adra1a^KO^) compared to AAV5-*gfaABC1D*::Cre-GFP-transduced *Adra1a^+/+^* (A-Adra1a^Cre-GFP^ controls, Fig.3I). Spontaneous Ca^2+^ transients were slightly reduced in amplitude in slices from A-Adra1a^KO^ compared to A-Adra1a^Cre-GFP^ controls (SupFig.5D,E). Remarkably, the large astrocyte Ca^2+^ elevation observed in response to NE in control conditions was completely missing in A-Adra1a^KO^ slices (Fig.3J,K), corroborating the functional ablation of astrocytic ɑ1A-ARs and the conclusion that NE-induced astrocyte Ca^2+^ responses result from the direct activation of ɑ1A-ARs on astrocytes (SupFig.2) (*12*). Strikingly, the effect of NE on fEPSP and PPF, while preserved in A-Adra1a^Cre-GFP^ control slices (Fig.3L), was also totally absent in slices taken from A-Adra1a^KO^ (Fig.3M-O). Therefore, collectively, our results allow the conclusion that NE modulates synaptic function not by signaling directly onto neurons but by engaging astrocyte activity (Fig.3P).

### NE acts by mobilizing ATP-adenosine signaling

An important corollary of the above conclusion is that the observed inhibitory effect of NE on synapses must be the result of an astrocyte output mobilized in response to NE. A variety of NE-driven astrocyte activities have been documented across brain regions, organisms, and preparations that are consequential to synaptic connectivity over seconds to minute timescales (*24*). Among them is the regulation of extracellular K^+^ (*49*), the supply of lactate (*50*), and the secretion of signaling molecules such as D-serine (*51*) and ATP (*22, 52*). The latter is of particular interest because, in the hippocampus, ATP is readily hydrolyzed into adenosine to act on the A1 receptor (A_1_R, Fig.4A), a Gi-coupled subtype of purinergic GPCR selective for adenosine. At CA3-CA1 synapses, A_1_Rs are predominantly presynaptic and their activation decreases transmitter release probability (*53*). Accordingly, direct applications of adenosine in hippocampal slices triggered an inhibitory effect on synaptic strength and presynaptic efficacy (*54*), the magnitude and timing of which strikingly resembled that of NE (SupFig.6A,B). Correspondingly, we found that blocking A_1_R with specific antagonists, CPT (200nM) or DPCPX (100nM), was sufficient to completely abolish the ability of NE to modulate synaptic transmission in slices from wild-type animals (Fig.4A-C and Sup Fig.6C). By contrast, inhibiting other purinergic receptors, including P_2_X and P_2_Y receptors, as well as adenosine A_2A_ and A_2B_ receptors, was without effect, pointing at a major role of adenosine rather than ATP and indicative of an A_1_R-specific mechanism (Fig.4A and SupFig.6D). To ascertain the presynaptic locus of A_1_R involved, *Adora1*^fl/fl^ mice were transduced with an AAV5-*hSyn*::Cre-GFP viral vector in the CA3 of the dorsal hippocampus (Fig.4D), which yielded robust GFP expression in 72% of CA3 neurons (i.e. pre-synaptic) and minimal expression in CA1 (6%, Fig.4E). Mice transduced with the same virus in the CA1 (i.e. post-synaptic neurons) were used as controls (Fig.4D) and showed a mirroring pattern of GFP expression (Fig.4E). In slices obtained 5 weeks later from CA3-injected mice, adenosine had a markedly reduced effect on fEPSPs and PPF compared to CA1-injected controls (Fig.4F,G, Table S1), confirming the loss of approximately 50% of presynaptic A_1_R function at CA3-CA1 synapses in these mice (CA3-specific *Adora1* knockdown, CA3-Ado1^KD^). Remarkably, while NE achieved a nominal inhibition of fEPSPs and accompanying PPF increase in slices from CA1-Ado1^KD^ mice, its effect was diminished by 59% in CA3-Ado1^KD^ slices (Fig.4H-J), consistent with an instrumental role of presynaptic A_1_Rs in NE neuromodulation of synapses. By contrast, inhibiting other modulators of presynaptic efficacy, such as metabotropic glutamate receptors (*55*), had no impact on the outcome of NE applications (Fig.4A & SupFig.6E).

**Fig. 4:**
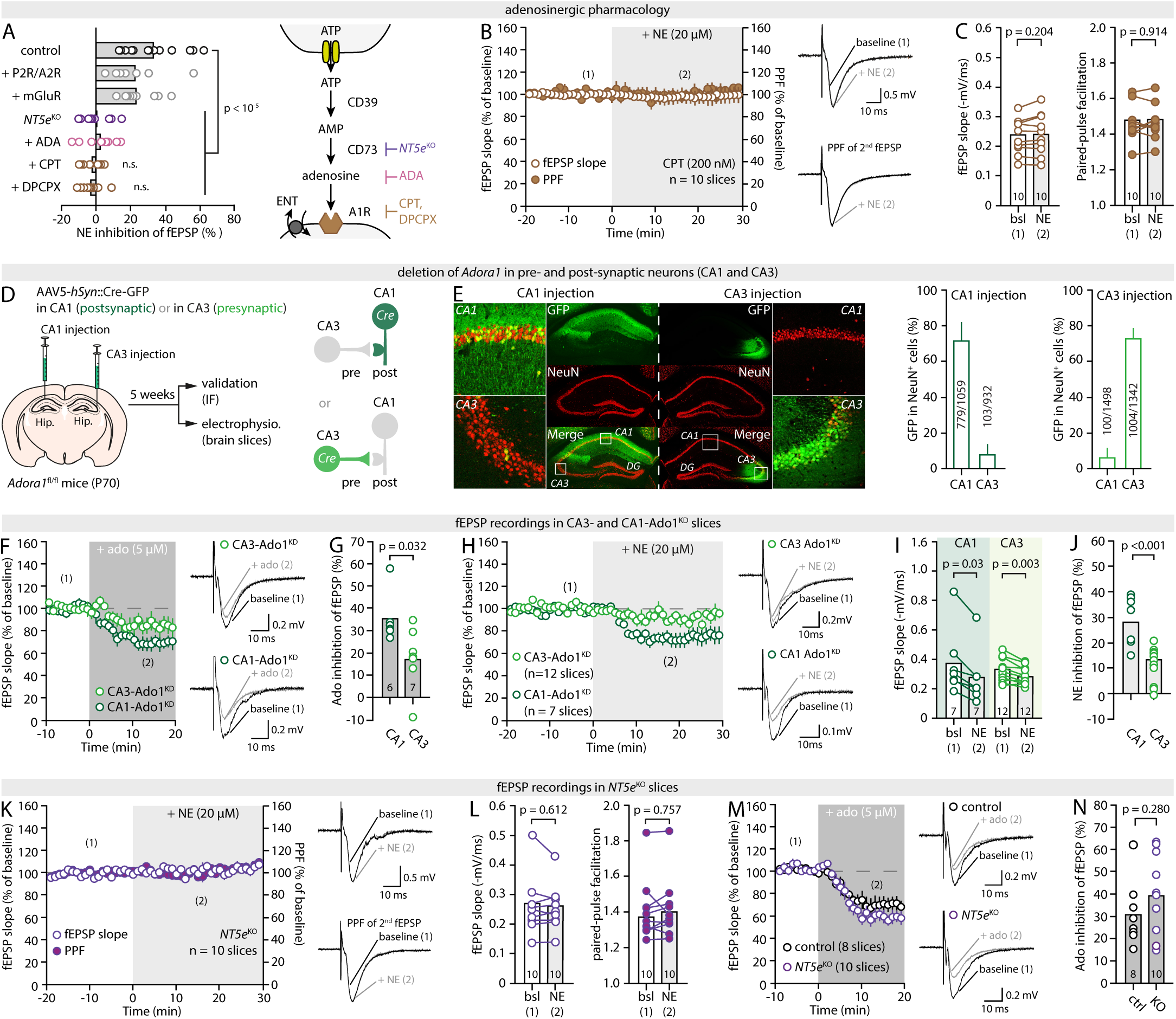
NE leverages ATP-Adenosine-A_1_R signaling to modulate synaptic efficacy. (**A**) *Left*, plot summarizing the effect of 20µM NE in the presence of A1R antagonists (CPT, 200nM, or DPCPX, 100nM), an adenosine scavenger (ADA, 1U/mL), a cocktail of P2X/P2Y (PPADS, 10µM), A_2A_ (ZM241385, 50nM) and A_2B_ (PSB603 50nM) receptor antagonists, a cocktail of mGluR inhibitors (CPPG, 5µM; MPEP, 3.6µM; YM298198, 2µM) or in slices from *NT5e*^KO^ mice. *Right*, schematic of the ATP-Adenosine-A_1_R pathways showing different points of genetic or pharmacological intervention. (**B**) Time-course of the effect of NE on fEPSPs and PPF in the presence of the A_1_R antagonist CPT and representative traces. (**C**) Pair-wise quantification for the experiments shown in (B). (**D**) Approach for the deletion of *Adora1* in CA3 or CA1 neurons. (**E**) Representative IHC images of Cre-GFP expression in CA1- and CA3-injected animals, along with quantification of regional specificity (n = 4 slices, 2 animals). (**F,G**) Time-course, representative traces and quantification of the effect of 5µM adenosine on fEPSPs in CA1-Ado1^KD^ and CA3-Ado1^KD^ slices. (**H-J**) Time-course, representative traces, pair-wise quantification and summary plot of the effect of 20µM NE on fEPSPs in CA1-Ado1^KD^ and CA3-Ado1^KD^ slices. (**K,L**) Time-course, representative traces and summary plot of the effect of 20µM NE on fEPSPs and PPF in *NT5e*^KO^ slices. (**M,N**) Time course, representative traces and quantification of the effect of adenosine on fEPSPs in *NT5e*^KO^ and control slices.

To interfere with adenosine directly, rather than A1Rs, we bathed slices in adenosine deaminase (ADA, 1U/mL, 15min prior and throughout the experiment), an enzyme that hydrolyzes extracellular adenosine into inosine, which is a by-product inoperant on A_1_R and A_2A/B_R at physiological concentrations (*56, 57*). ADA completely prevented the effect of NE on synapses (Fig.4A and SupFig.6G), confirming that adenosine is the effector that remodels synaptic function rather than NE itself. Seeking a pharmacology-free confirmation, we reasoned that NE should be ineffective in slices obtained from *NT5e*^KO^ mice lacking CD73, an ecto-5′-nucleotidase that catalyzes the conversion of interstitial AMP into adenosine, the last step in the enzymatic production of adenosine from extracellular ATP (Fig.4A). Hippocampal slices obtained from *NT5e*^KO^ mice appeared normal besides a notable hyperexcitability (not shown). Strikingly, NE caused no changes in fEPSP or PPF in *NT5e*^KO^ slices (Fig.4K,L). The lack of NE-induced effect was not due to the absence or downregulation of presynaptic A_1_Rs in these mice, or any overt compensations that would disrupt the basic synaptic machinery, because direct applications of adenosine elicited a normal fEPSP inhibition, accompanied by an increase in PPF, in the same slices (Fig.4.M,N and SupFig.6H,I). Of importance, this was also true for other conditions described above, including iβARK (SupFig.6F,J-L), CalEx (SupFig.3H,I,L), thapsigargin (SupFig.4E,F,I), and A-Adra1a^KO^ slices (SupFig5.F-H). In total, these results demonstrate that the ability of NE to remodel synaptic function relies fully on an ATP-adenosine-A_1_R-dependent control of presynaptic efficacy.

## Discussion

Our findings reveal that NE functionally remodels synapses by signaling through astrocytes. While presynaptic in nature, as documented in the past, the canonical effect of NE on synaptic function is unaffected by the deletion of its target receptor (ɑ1A-AR) on neurons, but abolished by silencing astrocytes, by suppressing astrocyte sensitivity to NE, and by genetic, enzymatic, or pharmacological interference at any level of an ATP-Adenosine-A_1_R pathway. Collectively, this fuels a model in which NE engages astrocyte Ca^2+^ dynamics, ATP/adenosine signaling, and the activation of pre-synaptic purinergic receptors to update synaptic weights in a non-Hebbian mechanism and reshape neuronal connectivity. As demonstrated in an accompanying paper by Chen et al. (*58*), an analogous astrocytic purinergic pathway recruited by NE is instrumental to behavioral state transition in the larval zebrafish, portraying astrocyte-based purinergic signaling as a general mechanism by which LC-NE activity remaps neural circuits. Consistently, seminal work in the fly larvae described astrocytes responsiveness to octopamine/tyrosine (Oct/Tyr) via the Oct/Tyr receptor, which are invertebrate analogs of NE and ɑ1-AR, respectively, and showed that it inhibits chemotaxis-regulating dopamine neurons via adenosine receptors (*22*). That astrocytes across three phylogenetically distant organisms share the same response to an evolutionarily conserved neuromodulator (NE/Oct/Tyr) indicates that they have evolved as an integral mechanism for monoaminergic systems to dynamically modulate brain circuits and behavior, and adds to mounting evidence that the role of astrocytes in brain function has been severely underestimated.

Importantly, past functional dissections of the effect NE at the synaptic and cellular level were conducted at a time when cell-specific and genetic approaches were unavailable or uncommon, and electron microscopy confirmations were scant or lacked systematic quantification. Additionally, alternative interpretations involving non-neuronal cells were given little credit because these cells were conceptualized as ‘inactive’ at the time. Hence, observations that NE elicited a presynaptic modulation sensitive to adrenergic pharmacology supported the conclusion that NE acted directly on presynaptic receptors. Yet, to our knowledge, there has been no formal demonstration that NE acts directly on synaptic terminals. In view of recent studies that paint a more diverse picture of adrenergic receptor expression and responsiveness across cell-types (*59*), our findings thus call for a systematic reexamination of other aspects of NE signaling, such as its effect on neuronal excitability.

More generally, our work supports a broad shift in how to conceptualize the cellular and molecular underpinnings of neuromodulation. The responsiveness of astrocytes to neuromodulators is not limited to NE but encompasses all canonical monoamines (*60, 61*) as well as acetylcholine (*62*) and oxytocin (*63*), which raises the question of the transferability of our findings. For instance, in the nucleus accumbens (NAc), dopamine recruits astrocyte Ca^2+^ activity and the subsequent release of ATP (*64*). Concomitantly, the motivational effect of dopamine has been linked to its inhibitory action on NAc synapses (*65*). Hence, the molecular rules by which all neuromodulators achieve their mesoscale and behavioral effects might benefit from being reevaluated with a more holistic perspective, for the mechanism we describe here might be generalizable to other neuromodulatory systems. Considering the importance of neuromodulation in conveying prediction error statistics in predictive coding theories of brain function (*9*), our findings pave the way to a better incorporation of astrocyte biology in systems neuroscience, towards multicellular models of brain function.

## Acknowledgments

Authors thank Dr. Robert W. Greene for his generous gift of the *Adora1*^f/f^ mice, Dr. Peter Bayguinov (assistant director of the Washington University Center for Cellular Imaging) and Dr. Mingjie Li (director of the WashU Hope Center Viral Vectors Core) for their technical assistance, and funding sources for their support. Authors we supported by the following funding sources: National Institutes of Health R01MH127163 (TP), R01DK128475 (VKS), R01NS102272 (JDD), Department of Defense grant W911NF-21-1-0312 (TP), The Brain & Behavior Research Foundation NARSAD Young Investigator Award 28616 (TP), The Whitehall Foundation grant 2020-08-35 (TP), The McDonnell Center for Cellular and Molecular Neurobiology Award 22-3930-26275U (TP).

## Author contributions

Conceptualization: TP

Methodology: KBL, YW, AY, TO, YZ, YD, SW, RM, PCS

Investigation: KBL, YW, AY, TO, YZ, YD, SW, RM, PCS

Visualization: TP

Funding acquisition: TP, JDD, VKS

Project administration: SW, RM, TP

Supervision: TP,

Writing – original draft: KBL, TP

Writing – review & editing: KBL, YW, YD, SW, JDD, VKS, TP

## Data availability

All data are available in the main text or the supplementary materials. All codes and methods for the STARDUST pipeline are currently available: https://doi.org/10.1101/2024.04.04.588196. All other data, code, and materials will be made available upon request. Adra1a flox/flox mice were obtained from Dr. Paul Simpson as part of materials transfer agreement A2022-1511 with UCSF. Adora1 flox/flox mice were obtained from Robert W Greene (UT Southwestern) without MTA. All other mouse lines and reagents are publicly available.

## Methods

### Animals

*Housing, breeding and genotyping*: All experiments were conducted in accordance with the guideline of the Institutional Animal Use Committee of Washington University in St. Louis School of Medicine (IACUC #20180184 and 21-0372, Animal Welfare Assurance #D16-00245). All mice were bred in our facility and group housed once weaned (2-5 per cage) except after surgical procedures when animals were singly housed until full recovery prior to being returned to home. All animals were kept on a 12-12 light-dark cycle (9am ON, 9pm OFF, or 11am ON, 11pm OFF) with access to food and water *ad libitum*. Adult male and female mice were used (P50-120, see below) and littermates of the same sex were randomly assigned to experimental groups. C57BL/6J mice (Stock #000664), *Dbh*^Cre^ mice (Stock #033951), and *NT5e*^KO^ mice (Stock #018986) were purchased from Jackson Laboratory. *Dbh*^Cre^ animals were maintained as hemizygous Cre while *NT5e*^KO^ mice were obtained and maintained as homozygous KO. *Adra1a*^fl/fl^ mice in which the first coding exon for ɑ1A-AR is flanked by loxP sites were generated (*8*) and contributed by Paul Simpson at the University of California, San Francisco, as part of a material transfer agreement (A2022-1511), and back-crossed with C57BL/6J mice. *Adora1*^fl/fl^ mice, in which the major coding exon for the A_1_R (homologous to human *Adora1* exon 6) is flanked with LoxP sites were generated (*66*) and generously gifted by Dr. Robert Green at the University of Texas Southwestern, backcrossed with C57BL/6J mice and bred as heterozygous. All genotyping primers are listed in Table S2. *Dbh*^Cre^ mice genotyping resulted in a 188bp and a 284bp band (Cre-positive) or a single 284bp product (Cre-negative). *NT5e*^KO^ mice genotyping yielded a 350bp product (compared to 235bp in C57BL/6J mice). *Adra1a*^fl/fl^ mice genotyping resulted in a ∼600bp product (floxed) and 524bp product (wildtype). *Adora1*^fl/fl^ mice genotyping resulted in a 302bp product (floxed) and/or a 268bp product (wildtype). All PCR was performed with an annealing temperature of 60°C, extension at 65°C, and denaturation at 95°C.

*Stereotaxic surgeries*: AAV micro-injections were carried out by intra-cranial stereotaxic surgeries (see AAV section for details on age, titration and volumes injected). Mice were anesthetized by 2-3% isoflurane inhalation. Buprenorphine Extended Release (1mg/kg, ZooPharm) was administered subcutaneously as a preoperative analgesic. Mice were placed on a Kopf stereotax and a small craniotomy was performed using a dental drill (Foredom). Micro-injections were bilateral and conducted with a micro-syringe pump controller (Kinder Scientific) via a 2µL Hamilton 7002 syringe (30 gauge) at a rate of 0.15 µL/min. 10min were allowed prior to removing the syringe. For GCaMP6f, CalEx, iβARK, and AAV-Cre injections targeting the CA1, coordinates were (in mm): -2.0 AP, +/-1.5 L, -1.5 D, relative to Bregma. For AAV-Cre injections targeting the CA3 (CA3-Ado1^KD^), coordinates were (in mm) -2.0 AP, +/- 2.2 L, -2.0 D. ChR2 injections in the LC of *Dbh*^Cre^ mice (P50) were performed at coordinates (in mm): -5.45 AP, +/- 1.25 L, -4.0 D. Incision site was sutured with 6-0 nylon sutures, and mice were given a 1mL warm saline subcutaneous injection and wet food, and monitored daily for 4 days. Once fully recovered, mice were returned to their home cage with their original littermates and allowed 3-12 weeks recovery.

### Electrophysiology

*Slice preparation and recording setup:* Electrophysiology experiments were carried out in acute hippocampal coronal slices (350 µm) obtained with a Leica VT1200s vibratome from adult male and female mice (P90-120) as previously described (*67*). After recovery (30min at 33°C and 45min at RT), slices were transferred to the recording chamber of a Scientifica SliceScope Pro 6000 system, where they were perfused with artificial cerebrospinal fluid (aCSF) saturated with 95%O_2_/5%CO_2_ at a flow rate of ∼1mL/min. The aCSF was maintained at 33°C (TC-344C Dual Channel Temperature Controller, Warner Instruments). The aCSF composition for slicing and recovery was (in mM) 125 NaCl, 3.2KCl, 1.25NaH_2_PO_4_, 26 NaHCO_3_, 1 CaCl_2_, 2 MgCl_2_, and 10 glucose (pH 7.3, 290-300 mOsm.L^-1^). The aCSF composition for recording was (in mM) 125 NaCl, 3.4KCl, 1.25NaH_2_PO_4_, 26 NaHCO_3_, 2 CaCl_2_, 1.3 MgCl_2_ and 10 glucose (pH 7.3, 290-300 mOsm.L^-1^).

*Field recordings*: Schaffer collaterals were electrically stimulated with a concentric tungsten electrode placed in the *stratum radiatum* of CA1, using paired stimulations (0.1 ms pulse, 200ms apart) continuously delivered at 0.05 Hz. Evoked field excitatory post-synaptic potentials (fEPSPs) were recorded using a glass electrode (2-5MΩ) filled with recording aCSF and placed in the *stratum radiatum* near the isosbestic point. Stimulation intensity (<150µA, 100µs) was set as needed to evoke a synaptic response without population spikes within the slope of the fEPSPs. Experiments were performed at 33° C in the presence of the GABA_A_ receptor blocker picrotoxin (50µM, Tocris). In experiments with viral expression, the recording and/or stimulating electrodes were placed in regions of high fluorescence to maximize the recruitment of, or recording from, synapses from transduced neurons or from synapses within the domain of transduced astrocytes. Data were acquired with a Multiclamp 700B amplifier (Molecular Devices) through a Digidata 1450A, sampled at 20kHz, filtered at 10kHz, and analyzed using pClamp11.0 software (Molecular Devices). Average traces were taken from 10-15min epochs before and after drug application at times indicated by numbers in parenthesis on time course graphs. The paired-pulse ratio was measured as the ratio of the slope of the second fEPSPs over the slope of the first fEPSP. Ratios greater than 1 (paired-pulse facilitation, PPF) are typical at CA3-CA1 synapses and reflect an enhanced glutamate release during the second synaptic response. This short-term potentiation is due to the incomplete Ca^2+^ buffering in pre-synaptic terminals during the short inter-pulse interval (200ms), leading to an additive effect on pre-synaptic free Ca^2+^ levels and an exaggerated vesicular release in response to the second pulse. Consequently, for a given inter-pulse interval, the extent of potentiation is inversely related to the vesicular release probability, i.e. a lower probability at baseline allows a greater short-term potentiation (greater PPF). By extension, a decrease in pre-synaptic release probability over the course of a recording will manifest as an increase in PPF. Changes in PPF were measured as PPF_change_ = (PPR_post_ – 1) / (PPR_pre_ – 1) and illustrated as “PPF of 2^nd^ fEPSP” on representative traces. These were obtained by vertically scaling full epoch traces (containing both the first and second fEPSPs) so the amplitude of first fEPSPs pre- and post-treatment would match, visually revealing differences (or absence thereof) in the extent to which the second fEPSP response were potentiated.

*Optogenetics:* 12 weeks after stereotaxic injection into the LC, hippocampal slices from ChR2-injected *Dbh*^Cre^ animals were obtained as above in low light settings. Slices were allowed to recover only for 10 minutes at 33°C, then at RT for at least one hour, protected from light as per previously described protocols (*38, 68*). For ChR2 excitation, square pulses of blue light (460nm, 5ms duration) were delivered through the 4X objective of an Olympus BX51WI microscope equipped with a CoolLED system. Illumination was applied to the entire field of view centered on the tip of the recording pipette and the output light power measured at the microscope objective was 15μW. A single 10min train of 1Hz light-pulses was delivered using a Master8 stimulus generator (AMPI). The area surrounding the recording electrode was examined for ChR2 expression after each recording (to avoid stimulating NE-release prior to recordings). Recordings with no noticeable EYFP-expression within the 40X field of view surrounding the tip of the recording electrode were excluded from analysis.

*Patch-clamp recordings:* Patch-clamp recording experiments were performed at 34°C using a heated recording chamber (ALA Scientific Instruments, HCS) and temperature controller (ALA, HCT-10) in slices obtained as above in the same conditions as those used for field recordings. Pyramidal CA1 neurons were identified in recording chamber in SliceScope Pro 6000 (Scientifica) under a 40X objective (NA 0.8, Olympus), using infrared DIC camera (Electro, Teledyne Photometrics). Patch-clamp recording pipettes were filled with a cesium gluconate solution containing (in mM): 130 Cs+-gluconate, 10 HEPES, 4 Mg-ATP, 0.5 Na-GTP, 5 Na-Phosphocreatine, 5 EGTA, 10 TEA-Cl (pH 7.3, 290 mOsm.l-1). Cells were patched in whole-cell configuration and voltage-clamp mode at -70mV, and access resistance (Ra) and holding current (Ih) were monitored throughout the experiment. Cells with Ra >25MΩ or Ih<−150pA at −70mV were discarded, as well as cells for which those parameters varied 20% or more during the recording. Stimulations of the Schaffer collaterals were continuously delivered at 0.33 Hz through a bipolar stimulating electrode (FHC #30202) positioned in the *stratum radiatum* near the patched cell (<200µm) via a stimulus isolator (WPI #A365). Stimulation intensity was set such as to elicit an evoked excitatory postsynaptic current (eEPSC) with ∼50% success rate. Data were acquired and recorded with a Multiclamp 700B amplifier (Molecular Devices.) through a Digidata 1550 (Molecular Devices), sampled at 10-20 kHz, filtered at 2 kHz, and analyzed using pClamp11 software (Molecular Devices.). The threshold used to determine successes vs failure (∼ -5pA) was usually 2 times the standard deviation of the baseline noise, determined from areas with no spontaneous activity on recorded epochs, and was verified by visual inspection and adjusted manually if needed. A response was considered a failure if its absolute peak amplitude was smaller than the threshold. Synaptic efficacy was defined as the rate of successful synaptic events, synaptic potency as the mean peak amplitude of successful events (i.e., excluding failures) and synaptic strength as the mean peak amplitude of all events (failures + successes), over a defined epoch.

### Calcium imaging

GCaMP6f-based recordings of calcium activity in *stratum radiatum* astrocytes were performed in acute hippocampal slices obtained as above from C57BL/6J mice microinjected with AAV5-*gfa*ABC1D::lck-GCaMP6f in the CA1 *stratum radiatum* 3-4 weeks prior. Two-photon laser scanning microscopy (2-PLSM) images of GCaMP6f fluorescence (field of view 302×302µm, resolution 512×512 pixels) were taken at a 1Hz frame rate with a Nikon A1RHD25 MP microscope (920nm excitation laser, 515/530 filter, 25x objective, 1.1 numerical aperture submersion lens, 1.6x optical zoom). The aCSF (composition as above, perfusion rate ∼2.5mL/min) contained tetrodotoxin (1µM) to prevent neuronal activity. Recordings consisted of a 10min baseline, during which spontaneous Ca^2+^ activity was captured, immediately followed by the perfusion of NE and continuous recording for an additional 10min. Images of mCherry or RFP-reporter expression (in CalEx, iβARK, and A-Adra^KO^ slices) were taken with a 605/615 emission filter for *ad-hoc* cell segmentation.

*Calcium activity analysis:* 2-PLSM recordings were saved as TIF files and analyzed with the custom-made Python pipeline for astrocyte Ca^2+^ analysis, STARDUST (Spatio-Temporal Analysis of Regional Dynamics & Unbiased Sorting of Transients (*69*)). Briefly, an output map of regions of activity (ROAs) was obtained from the temporal projection of active pixels and overlaid on raw fluorescence recordings in ImageJ. The resulting regionalized time-series fluorescence data were fed into the STARDUST Python pipeline. This yielded normalized time series of the signal intensity (ΔF/F_0_) for each ROA, from which Ca^2+^ transient were automatically identified (3SD threshold above baseline) and their properties (amplitude, frequency, kinetics) evaluated using classic criteria (*69*). Spontaneous activity data were averaged across ROAs for each cell and presented as ‘averaged ROA” data per astrocyte (<ROAS>/astrocyte). NE-induced Ca^2+^ time-series were obtained from entire cells by segmenting astrocytes in ImageJ to generate a cell mask. Like for spontaneous activity measurements, the mask and the raw fluorescence data were then fed into the STARDUST pipeline for signal and feature extraction.

### Reagents

*Drugs:* Adenosine, YM-298190, 8-cyclopentyl-1 3-dimethylxanthine (CPT), and adenosine deaminase (ADA), were purchased from MilliporeSigma. Adenosine and YM-298190 were dissolved in DI water and stored at -20°C. CPT was dissolved in DMSO and stored at -20°C. Adenosine deaminase was stored at 4°C. LY746-314 was purchased from Santa Cruz, dissolved in DMSO, and stored at -20°C. All other drugs were purchased from Tocris Biosciences, dissolved in DI water (norepinephrine, propranolol, A61603, PPADS, NBQX, and D-AP5), DMSO (yohimbine, prazosin, silodosin, PSB603, ZM 241385, MPEP, and SCH442416), ethanol (picrotoxin and DPCPX), 1eqNaOH (CPPG), or citric acid (tetrodotoxin) and stored at -20°C.

*Adeno-associated viruses:* AAV5-*gfaABC1D*::lckGCaMP6f (Addgene #52924-AAV5) was co-injected at a 3:1 ratio with AAV5-*gfaABC1D*::mCherry (Addgene #58909-AAV5, 1µL with a total of 1×10^10^ genome copies injected), or at a 1:1 ratio with AAV5-*gfaABC1D*::hPMCA2-mCherry (Addgene #111568, packaged by the Washington University Hope Center Viral Vector Core, 2µL, 2×10^10^ genome copies total injected) or AAV5-*gfaABC1D*::iβARK-p2A-mCherry (Addgene #117691, packaged by the Hope Center Viral Vector Core2µL, total 2×10^10^ genome copies injected). Micro-injections were done bilaterally in P70-80 in C57Bl/6J male and female mice. For electrophysiology experiments, AAV5-*gfaABC1D*::hPMCA2-mCherry and AAV5-*gfaABC1D*::iβARK-p2A-mCherry were injected alone (1×10^10^ genome copies injected, 1µL) in C57BL/6J male and female mice (P70-80).

AAV5-*gfaABC1D*::Cre-GFP (Vector Biolabs #VB1131) or AAV5-*hSyn*::Cre-GFP (Addgene #100896-AAV5) were injected in *Adra1a*^fl/fl^ mice and *Adra1a*^+/+^ littermates (P70-80) at 1×10^10^ genome copies (1µL). In a subset of experiments *Adra1a*^fl/fl^ mice and *Adra1a*^+/+^ littermates were co-injected with AAV5-*gfaABC1D*::lckGCaMP6f and AAV5-*gfaABC1D*::Cre-mCherry (Vector Biolabs #VB1132, 1:1 ratio, 2µL, 2×10^10^ genome copies total injected). AAV5-*hSyn*::Cre-GFP was also injected in *Adora1*^fl/fl^ animals (P60) at 3×10^9^ genome copies (0.3 µL) to generate CA1- and CA3-specific Ado1^KD^.

AAV5-*EF1α*::DIO-hChR2(H134R)-EYFP (Addgene, 20298-AAV5) or AAV9-*synP::*DIO-EGFP (Addgene 100043-AAV9) were injected at 3×10^9^ genome copies (0.25 µL) in *Dbh*^Cre^ mice (P50). All reported injections reflect the volume and viral genome copies injected into each hemisphere. All injections were bilateral.

***FACS and RT-PCR:*** A- and N-Adra1a^KO^ animals were deeply anesthetized under isoflurane anesthesia, rapidly decapitated and hippocampi were dissected in ice-cold PBS (Fisher, BP39950). The tissue was dissociated into a single cell suspension and the debris were removed using the Adult Brain Dissociation Kit (Miltenyi Biotec, 130-107-677) and gentleMACS program 37C_ABDK_02 according to manufacturer’s instructions. The cells were re-suspended in FACS buffer (PBS, 1% BSA) to a target concentration of 1-5M cells/mL and were filtered through a 70 µm cell strainer prior to sorting for GFP-positive cells. All samples were maintained on ice until sorted or set aside as “bulk” samples to be digested without sorting. Samples were loaded into the Cytoflex SRT Cell Sorter (Beckman Coulter) equipped with a 100 µm nozzle and a 488 nm laser with a 525/40 bandpass filter to collect GFP fluorescence. FSC and SSC gates and laser gains were determined using GFP-negative cells from non-injected animals. A two-step gating strategy was used to identify and remove doublets. First, a FSC-A x SSC-A gate was used to select cells and remove debris. Then, a pulse geometry gate (FSC-A x FSC-H) was used to select singlets. A final gate for GFP+ and GFP-cells were used to select the cell populations to sort. The cells were sorted into 5mL round-bottom tubes containing cold FACS buffer. After sorting, the cells were spun at 500 xg for 5 min at 4°C in a swinging bucket rotor and the supernatant was aspirated. To harvest genomic DNA, the cells were re-suspended in lysis buffer (25mM NaOH, 0.2mM EDTA, pH 12) and heated to 95° C for 30 min. Equi-volume neutralization buffer (40mM Tris-HCl pH 5) was added to each sample and the gDNA was stored at 4°C. RT-PCR was performed using the *Adra1a* primers (Table S2) to assess presence of the unmodified *Adra1a* gene and separately of *Actb* (encoding β-Actin, Table S2). Gels were then imaged using a ChemiDoc MP Imaging System (Biorad, 1s exposure). Band intensity was then calculated in ImageJ. *Adra1a* band intensity was normalized to β-Actin for each sample. Loss of *Adra1a* was assessed as a normalized ratio of *Adra1a*/*Actb* for each sorted and bulk sample.

***Immunofluorescence and confocal imaging***: Mice were deeply anesthetized with isoflurane inhalation anesthesia, and transcardially perfused with ice cold PBS followed by 4% PFA (Thermo Scientific). Brains were then removed from the skull and postfixed in 4% PFA for 24 hours (4°C), followed by 30% sucrose for 2 days at 4°C. 50µm coronal free-floating sections were obtained on a vibratome (Leica VT1000) in cold PBS and stored at 4°C in PBS for up to one week. Sections were washed and permeabilized in 0.5% PBS-TritonX-100 for 10min, then washed three times for 5min per wash in PBS. Sections were then blocked in 5% normal goat serum (NGS, Abcam, AB7481) in 0.5% PBS-T for 30min. Sections were incubated with primary antibodies in 1% NGS in 0.5% PBS-T at 4°C overnight on an orbital shaker (gentle rotation), then washed in PBS for 5min three times and incubated with secondary antibodies in 1% NGS in 0.5% PBS-T for two hours at RT, protected from light. The list of primary (Table S3) and secondary (Table S4) antibodies is provided below. Sections were finally washed in PBS, mounted on Superfrost plus slides (VWR, 12-550-15), and sealed with Vectashield hardset mounting media with DAPI (Vector Laboratories, H-1800-10). Confocal images were acquired with a 1246 x 1246 resolution on a Zeiss Airyscan 800 with a 20X lens (NA=0.8) for z-stacks, or 10X (NA=0.45) for tiled images. For all images, the following laser lines were used (in nm): 405, 488, 561, 633. Images were then processed in Fiji for cell counting as described (*70*). Specificity was determined as the percentage of all reporter-expressing cells that are also positive for a marker of the desired cell-type (e.g., % of GFP-positive cells also positive for GFAP), while efficiency was measured as the percent of targeted cells that also express the reporter (e.g., % of GFAP-positive cells that are also GFP positive).

***Quantification and statistical analysis:*** Statistical analysis was conducted using Graphpad Prism 10 softwar**e (**GPS-1564787-LIRT-FEC03). For the pair-wise comparisons for the effect of drugs within an experiment, paired Student’s *t*-test were performed after validating normality (using a Shapiro–Wilk test). The effect of drugs across multiple conditions was analyzed using one-way ANOVA followed by Tukey’s post-hoc test for multiple comparisons. A Pearson’s test was used to assess the correlation of the inhibitory effect of NE and the change in PPF (Fig.1) and between the inhibition by NE and loss of astrocyte Ca^2+^ response across astrocyte silencing conditions (Fig.2). For Ca^2+^ imaging data, datasets were analyzed with permutation tests. For validation of the loss of *Adra1a* in A- and N-Adra^KO^ mice, an unpaired Student’s *t*-test was used. All data are shown as average ± s.e.m. (standard error to the mean) or average and individual data points. All statistical data are available in Table S1.

## Supplementary

**SupFig. 1:**
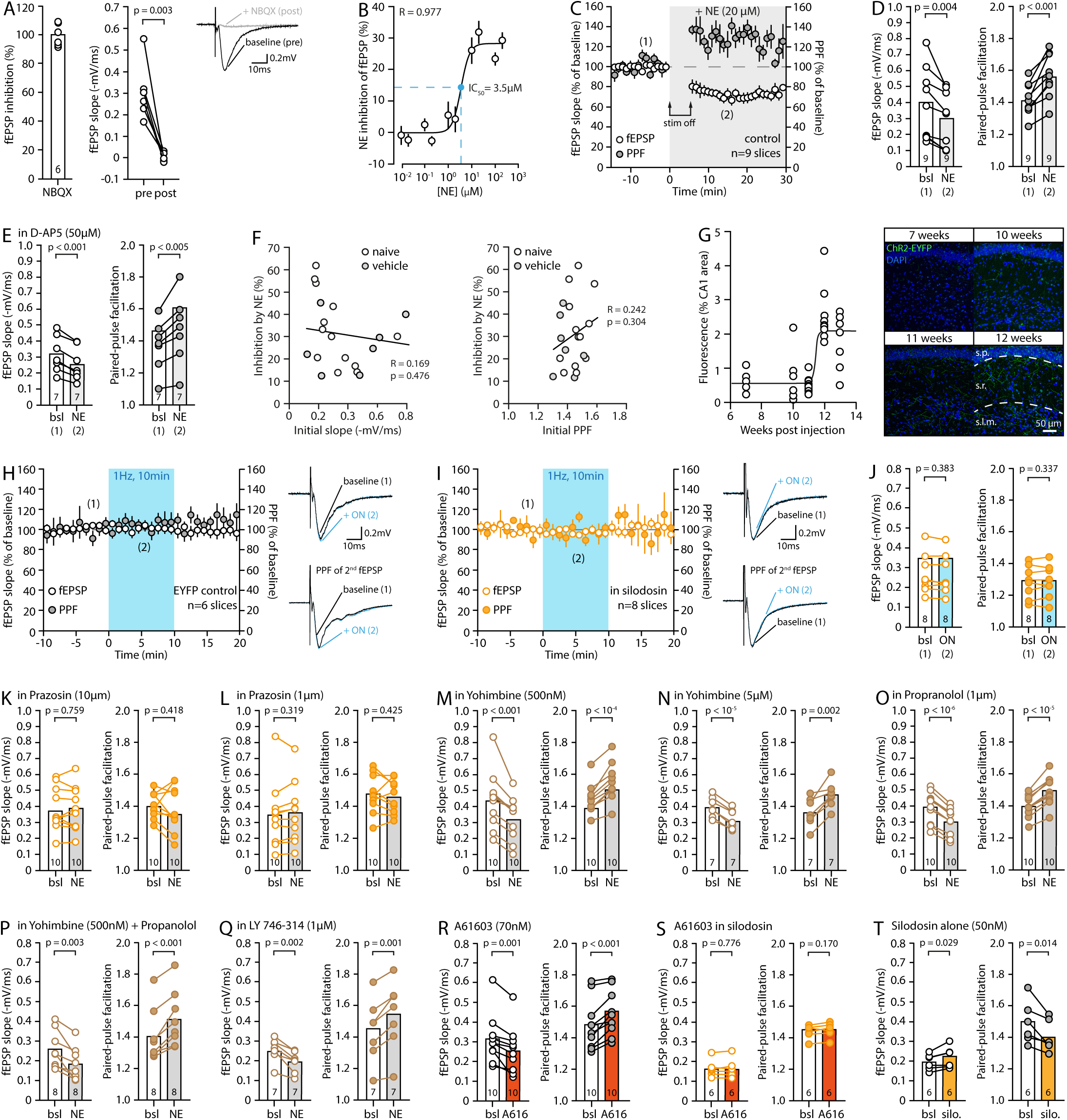
Effect of NE on the AMPAR-mediated fEPSPs. **(A)** Pairwise and summary quantification of the effect of the AMPAR antagonist NBQX (10µM) on fEPSPs and representative traces. **(B)** Dose-response curve of the inhibitory effect of 20µM NE on fEPSP slope (n = 5 to 7 slices per concentration). **(C,D)** Time course and pairwise quantification of the effect of NE on fEPSP and PPF with stimulations of Schaffer collaterals paused for five minutes at the onset of NE application. **(E)** Pairwise quantification of the effect of 20µM NE on fEPSP and PPF in the presence of the NMDAR antagonist D-AP5 (50µM). **(F)** Plots of the inhibitory effect of NE as a function and the initial fEPSPs slope (left) or initial PPF (right) across control conditions. The naïve and vehicle condition (DMSO, 0.01%) are denoted separately, but the linear regression, correlation coefficient and p-values are shown for the control condition as a whole. **(G)** Quantification and representative images of ChR2-eYFP expression of LC-NE projections in the hippocampal CA1 at 7, 10, 11, 12 and 13 weeks (n = 6-12 slices, from 1-2 animals). **(H)** Time course and representative traces showing the effect of the optical stimulation of LC-NE fibers on fEPSPs and PPF in EYFP-control slices. **(I,J)** Time course, representative traces and pairwise quantification showing the effect of the optical stimulation of LC-NE fibers on fEPSPs and PPF in the presence of silodosin (50nM). **(K-Q)** Pairwise quantification of the effect of 20 µM NE on fEPSP slope and PPF in prazosin (α1-AR antagonist, (K,L)), yohimbine (α2-AR antagonist, (M,N)), propranolol (β antagonist, (O)), yohimbine and propranolol (P), and LY746-314 (α1B-AR antagonist, (Q)) at indicated concentrations. **(R,S)** Pairwise quantification showing the inhibitory effect of the α1A-AR agonist A61603 (70nM) on fEPSP and PPF (R) and its blocked by silodosin (S). **(T)** Pairwise quantification showing the potentiating effect of silodosin perfusion (50nM) on fEPSP and inhibitory effect on PPF.

**SupFig. 2:**
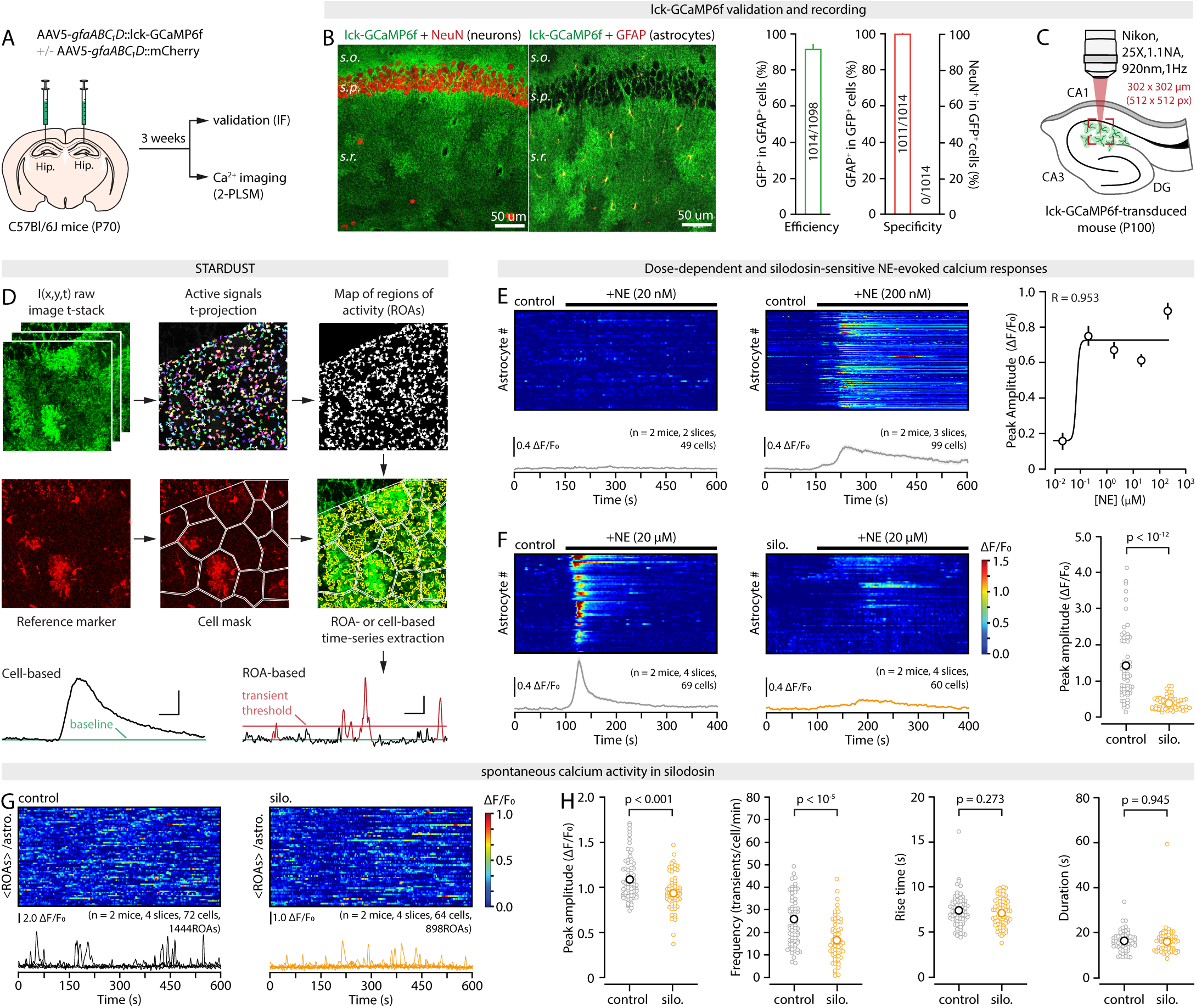
Imaging and analysis of astrocyte Ca^2+^ responses to NE and α1A-AR pharmacology. **(A)** Schematic of micro-injections for lck-GCaMP6f expression in astrocytes in the *s. radiatum* of the CA1. **(B)** IHC of lck-GCaMP6f expression in CA1 astrocytes and quantification of efficiency and specificity. **(C)** Schematic of the 2-PSLM conditions for astrocyte lck-GCaMP6f imaging in hippocampal slices. **(D)** Overview of the STARDUST analysis workflow for ROA and cell-based timeseries analysis. **(E)** *Left,* Kymographs of whole astrocyte Ca^2+^ signals (each row represents a single cell) and average ΔF/F_0_ traces (± s.e.m.) across all astrocytes, in responses to the application of NE at indicated concentrations. *Right*, Dose response curve showing the peak amplitude of the astrocyte Ca^2+^ response as a function of NE concentration. **(F)** *Left,* Kymographs of whole astrocyte Ca^2+^ signals (each row represents a single cell) and average ΔF/F_0_ traces (± s.e.m.) across all astrocytes, in response to the application of 20µM NE in control conditions and in the presence of silodosin (50nM, α1A-AR antagonist). *Right*, Quantification of the peak astrocyte Ca^2+^ response to 20µM NE in control and silodosin. **(G)** Kymographs (each row shows the average fluorescence across ROAs of a single astrocyte) and 5 representative ΔF/F_0_ traces (from individual ROAs) showing spontaneous astrocyte Ca^2+^ transients in control and silodosin conditions. **(H)** Quantification of the peak amplitude, frequency, and kinetics of spontaneous astrocyte Ca^2+^ transients in control conditions and in the presence of silodosin.

**SupFig. 3:**
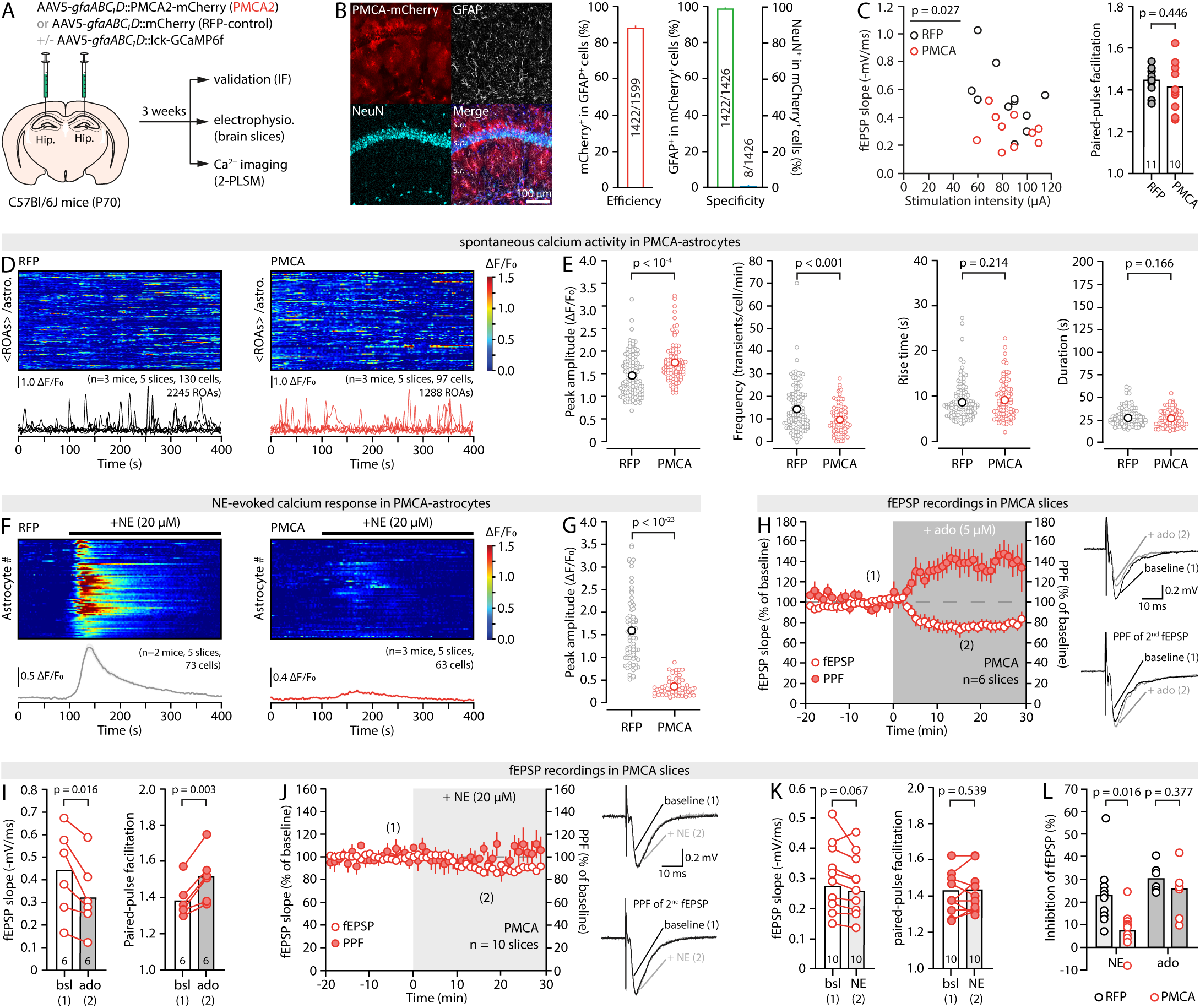
CalEx blocks the effect of NE on astrocyte Ca^2+^ and synapses. **(A)** Schematic micro-injections to express the CalEx actuator PMCA2 (plasma membrane Ca^2+^ pump) in CA1 astrocytes. **(B)** IHC of PMCA2 expression in CA1 *s. radiatum* astrocytes and quantification of specificity and efficiency. **(C)** Plot of the stimulation intensity/fEPSP slope relationship (left, unpaired Student’s *t*-test on slope/stim ratio) and summary bar graphs of PPF values (right) in RFP-control and CalEx slices at baseline. **(D,E)** Kymograph (each row shows the average fluorescence across ROAs of a single astrocyte), 5 representative ΔF/F_0_ traces (from individual ROAs), and quantification of the peak amplitude, frequency, and kinetics of spontaneous Ca^2+^ transients in RFP-control and CalEx slices. **(F,G)** Kymographs (each row represents a single cell), average ΔF/F_0_ traces (± s.e.m.) across all astrocytes, and quantification of the peak amplitude, in response to 20µM NE application in RFP-control and CalEx slices. **(H)** Time course and representative traces of the effect of adenosine on fEPSP and PPF in CalEx slices. **(I)** Pairwise quantification of the effect of adenosine on fEPSP and PPF in CalEx slices. **(J,K)** Time course, representative traces, and pairwise quantifications of the effect of NE on fEPSP and PPF in CalEx slices. **(L)** Summary bar graphs showing the inhibitory effect of NE and adenosine on fEPSP in RFP-control and CalEx slices. The p-values in (L) are from ANOVAs across iβark, CalEx, and RFP-control conditions followed by Tukey’s *post-hoc* test, reflecting the fact that the RFP-control condition for the experiments shown in (L) is the same across iβark (Fig.2) and CalEx (this figure) experiments.

**SupFig. 4:**
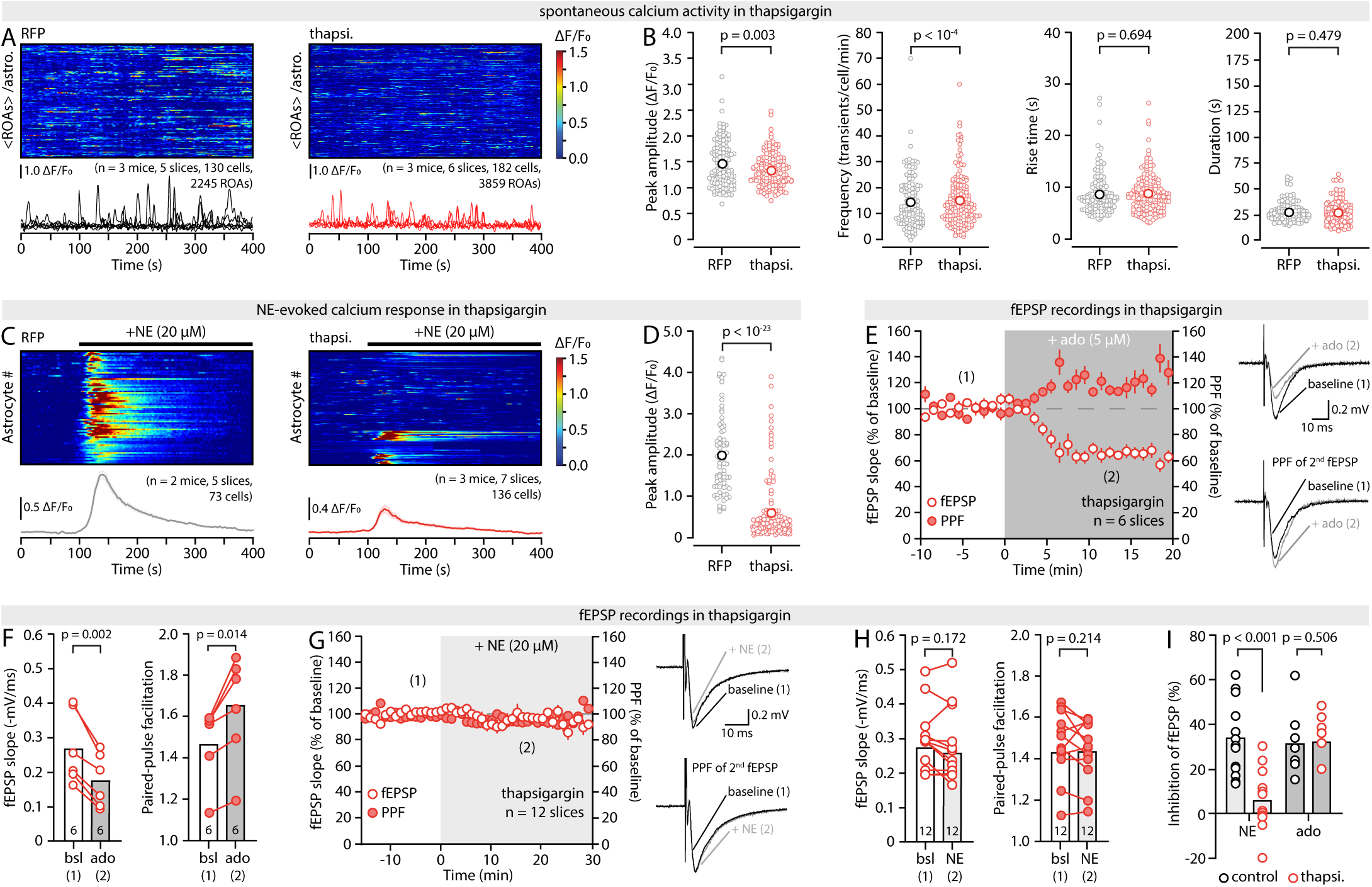
Thapsigargin blocks the effect of NE on astrocyte Ca^2+^ and synapses. **(A,B)** Kymograph (each row shows the average fluorescence across ROAs of a single astrocyte), 5 representative ΔF/F_0_ traces (from individual ROAs), and quantification of the peak amplitude, frequency, and kinetics of spontaneous Ca^2+^ transients in TTX alone (RFP-control) and TTX + thapsigargin conditions. All slices were obtained from animals with RFP-transduced astrocytes for cell-segmentation purposes. Thapsigargin was bath applied 20min prior to the start of recording. **(C,D)** Kymographs (each row represents a single cell), average ΔF/F_0_ traces (± s.e.m.) across all astrocytes, and quantification of the peak amplitude, in response to 20µM NE application in RFP-control and thapsigargin conditions. **(E,F)** Time course, representative traces, and pairwise quantification of the effect of 5µM adenosine on fEPSP and PPF in thapsigargin-treated slices. **(G,H)** Time course, representative traces, and pairwise quantifications of the effect of 20µM NE on fEPSP and PPF in thapsigargin-treated slices. **(L)** Summary of the inhibitory effect of NE and adenosine on fEPSP in control and thapsigargin-treated slices.

**SupFig. 5:**
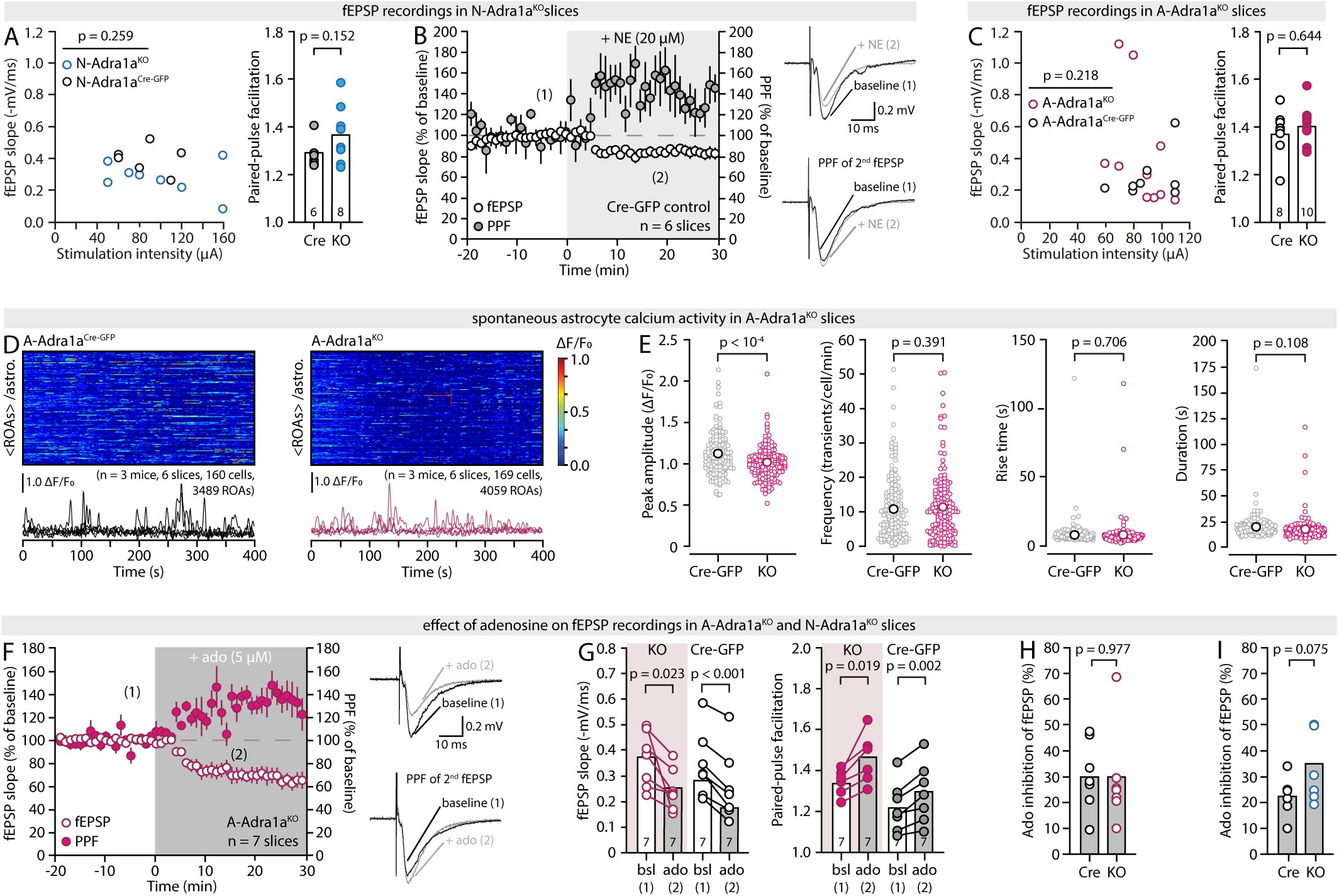
Ca^2+^ and synaptic recordings in A- and N-Adra^KO^ slices. **(A)** Plot of the stimulation intensity/fEPSP slope relationship (left, unpaired Student’s *t*-test on slope/stim ratio) and summary bar graphs of PPF values (right) in N-Adra^KO^ and N-Adra^Cre-GFP^ slices at baseline. **(B)** Time course and representative traces of the effect of 20µM NE on fEPSP and PPF in N-Adra^Cre-GFP^ slices. **(C)** Plot of the stimulation intensity/fEPSP slope relationship (left, unpaired Student’s *t*-test on slope/stim ratio) and summary bar graphs of PPF values (right) in A-Adra^KO^ and A-Adra^Cre-GFP^ slices at baseline. **(D,E)** Kymograph (each row shows the average fluorescence across ROAs of a single astrocyte), 5 representative ΔF/F_0_ traces (from individual ROAs), and quantification of the peak amplitude, frequency, and kinetics of spontaneous Ca^2+^ transients in A-Adra^KO^ and A-Adra^Cre-GFP^ slices. **(F)** Time course and representative traces of the effect of 5µM adenosine on fEPSP and PPF in A-Adra^KO^ slices. **(G)** Pairwise quantification of the effect of 5µM adenosine on fEPSP and PPF in A-Adra^KO^ and A-Adra^Cre-GFP^ slices. **(H)** Summary plot of the inhibitory effect of adenosine on fEPSP in A-Adra^KO^ and A-Adra^Cre-GFP^ slices. **(I)** Summary plot of the inhibitory effect of adenosine on fEPSP in N-Adra^KO^ and N-Adra^Cre-GFP^ slices.

**SupFig. 6:**
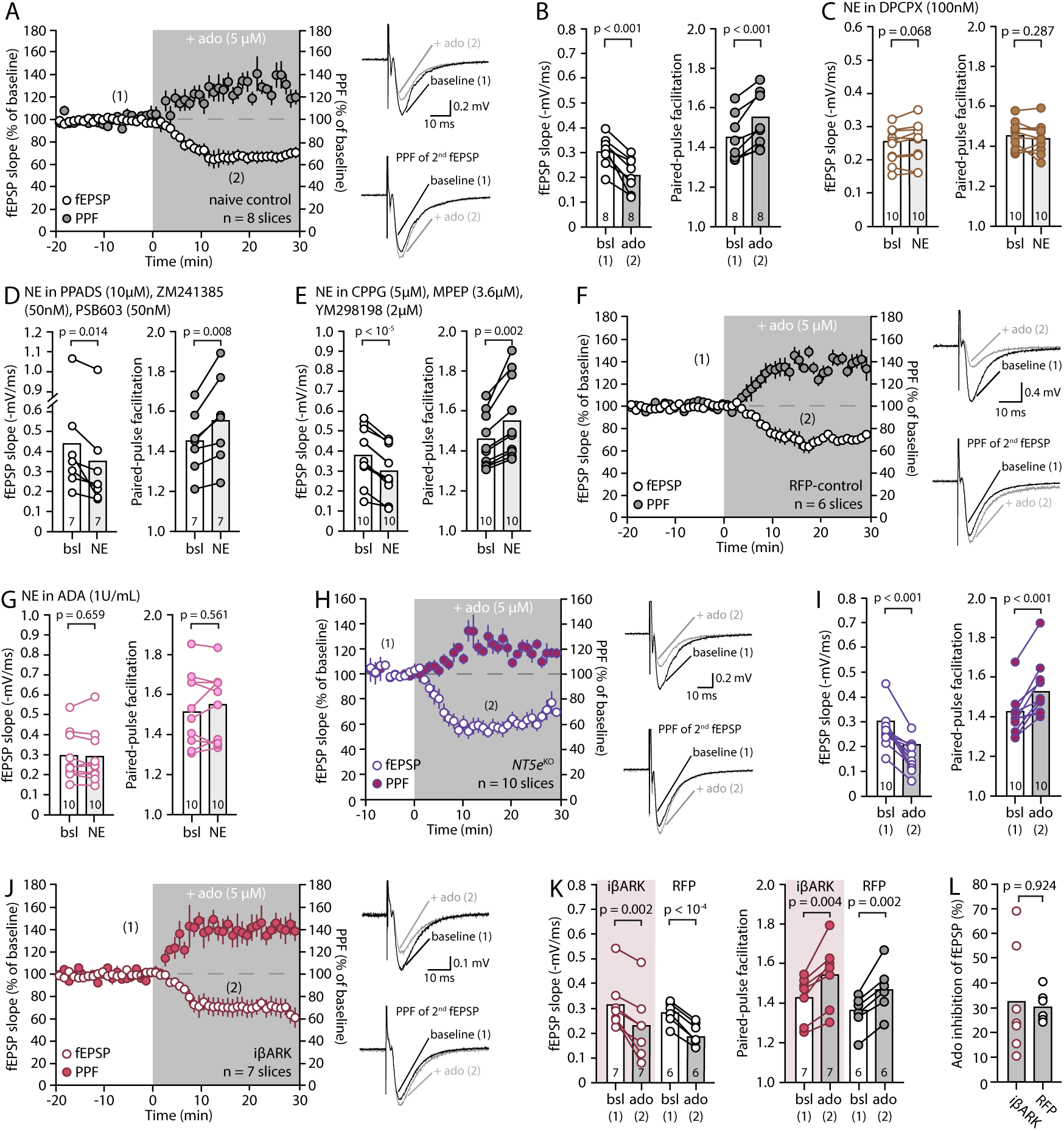
Adenosine signaling is central to the effect of NE and preserved in NT5e^KO^ animals and across astrocyte Ca^2+^ interventions. **(A,B)** Time course, representative traces, and pairwise quantifications of the effect of adenosine on fEPSP and PPF in slices from naïve animals. **(C-E)** Pairwise quantification of the effect of 20µM NE on fEPSP and PPF in DPCPX (100nM, C), a cocktail of P2X and P2Y inhibitors (D), and a cocktail of mGluR inhibitors (E). **(F)** Time courses and representative traces of the effect of 5µM adenosine on fEPSP and PPF in RFP-control slices. **(G)** Pairwise quantification of the effect of 20µM NE on fEPSP and PPF in ADA (1U/mL). **(H,I)** Time course, representative traces, and pairwise quantification of the effect of 5µM adenosine on fEPSP and PPF in NT5e^KO^ slices (paired Student’s *t*-tests). **(J)** Time courses and representative traces of the effect of 5µM adenosine on fEPSP and PPF in iβARK slices. (**K**) Pairwise quantification of the effect of adenosine on fEPSP and PPF in iβARK and RFP-control slices **(L)** Summary plot of the inhibitory effect of adenosine on fEPSP in iβARK and RFP-control slices. The p-value in (L) is from an ANOVA across iβark, CalEx and RFP-control conditions followed by Tukey’s *post-hoc* test, reflecting the fact that the RFP-control condition for fEPSP recordings experiments is common to CalEx (SupFig.3) and iβark (Fig.2).

**Table S1:**
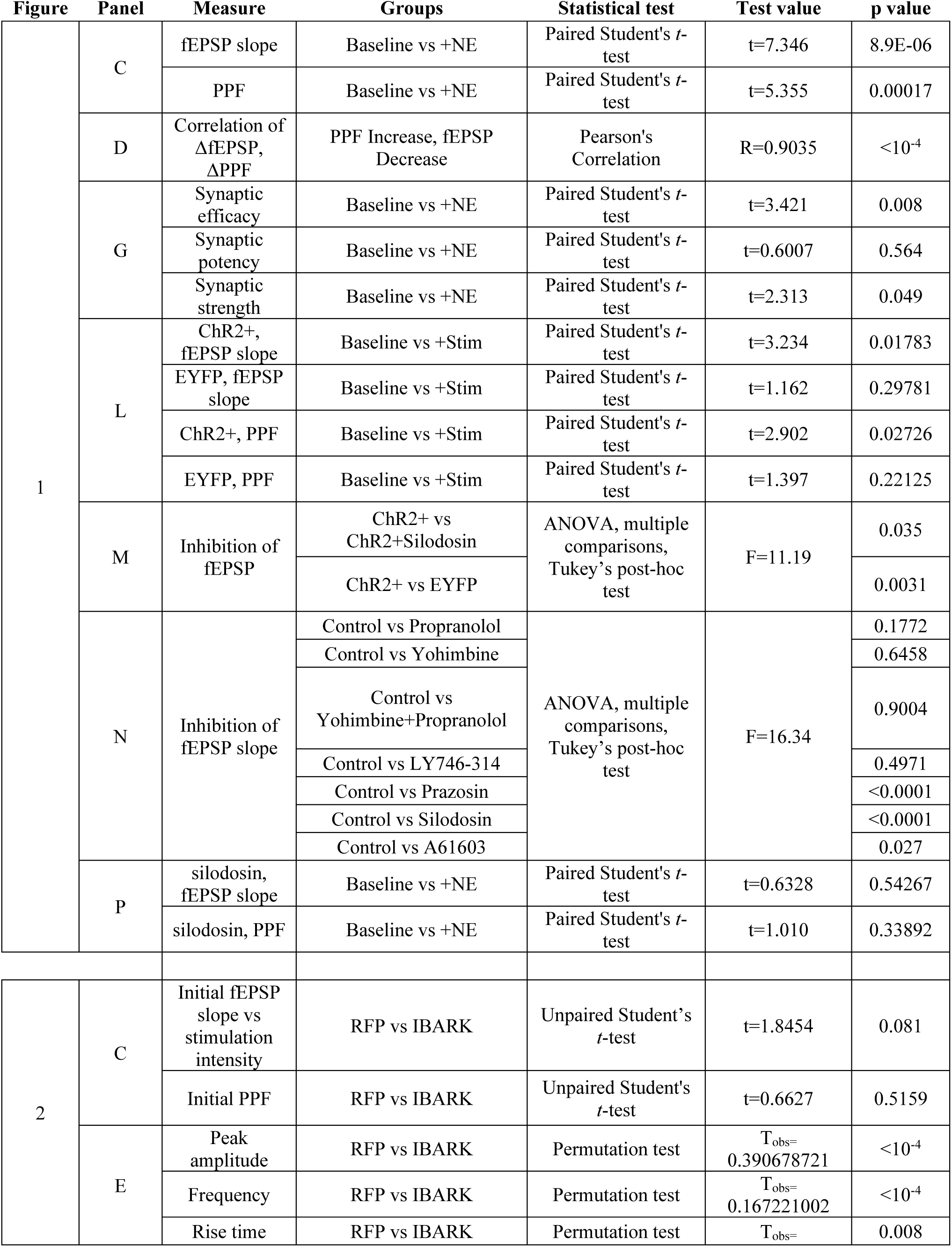
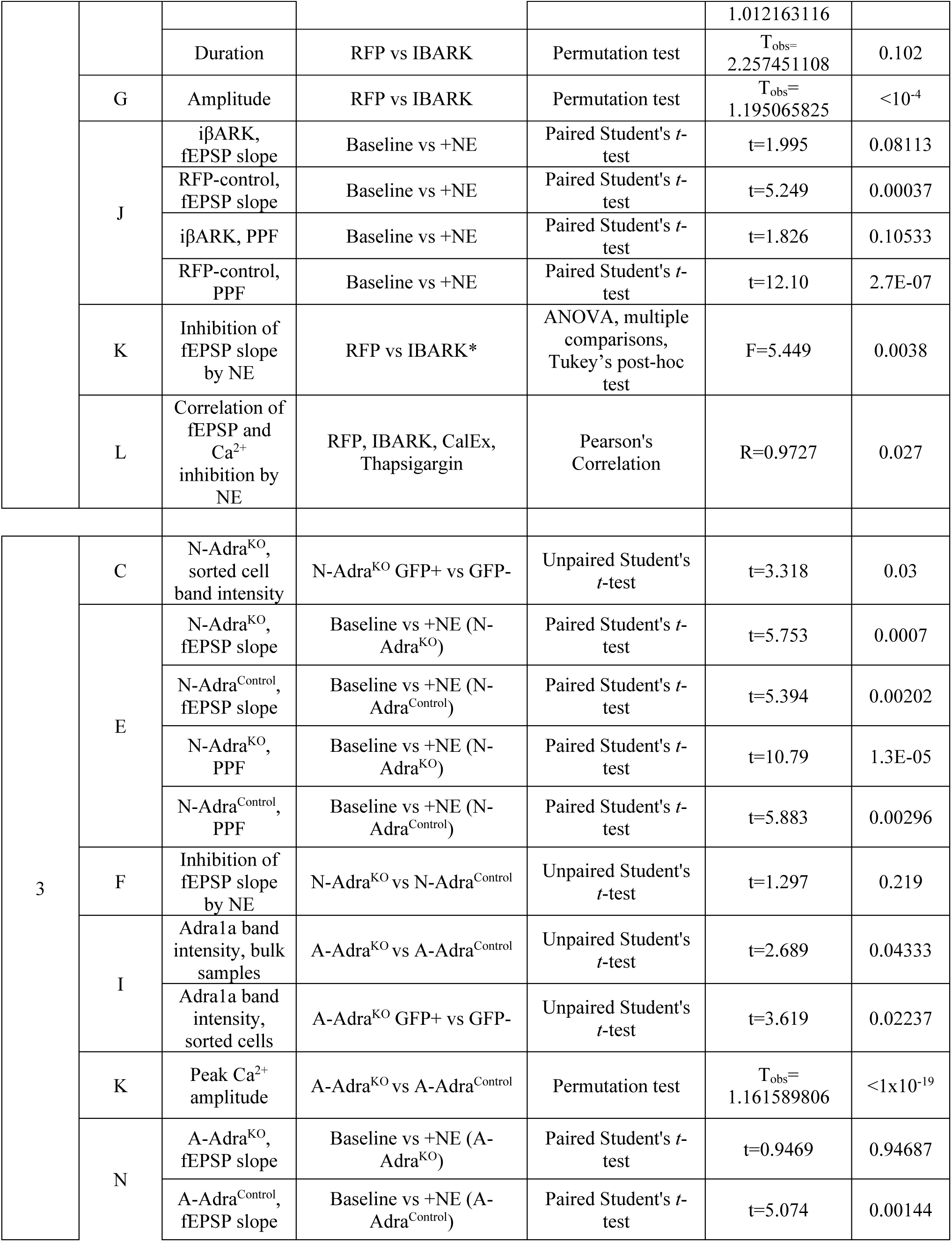
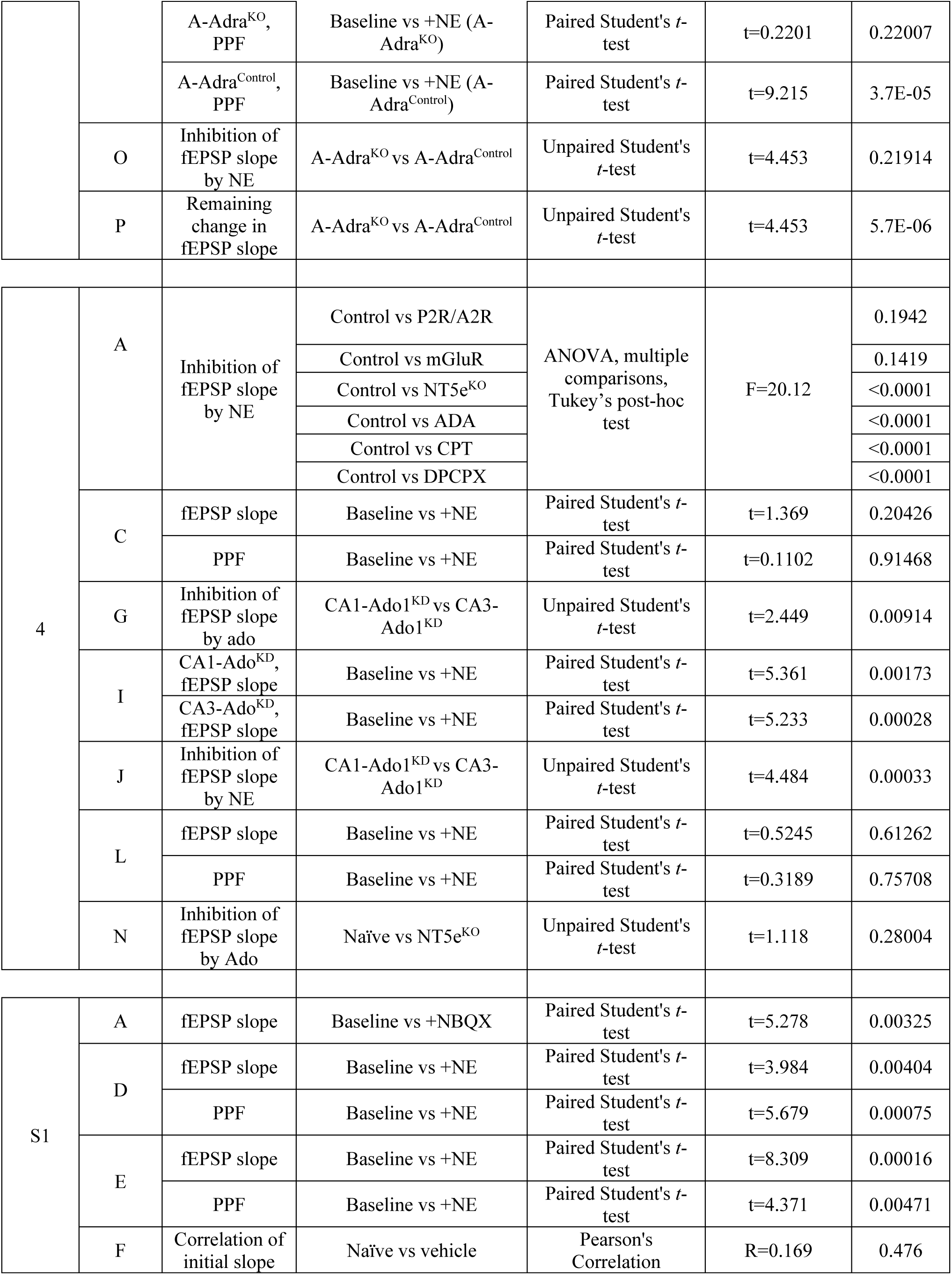
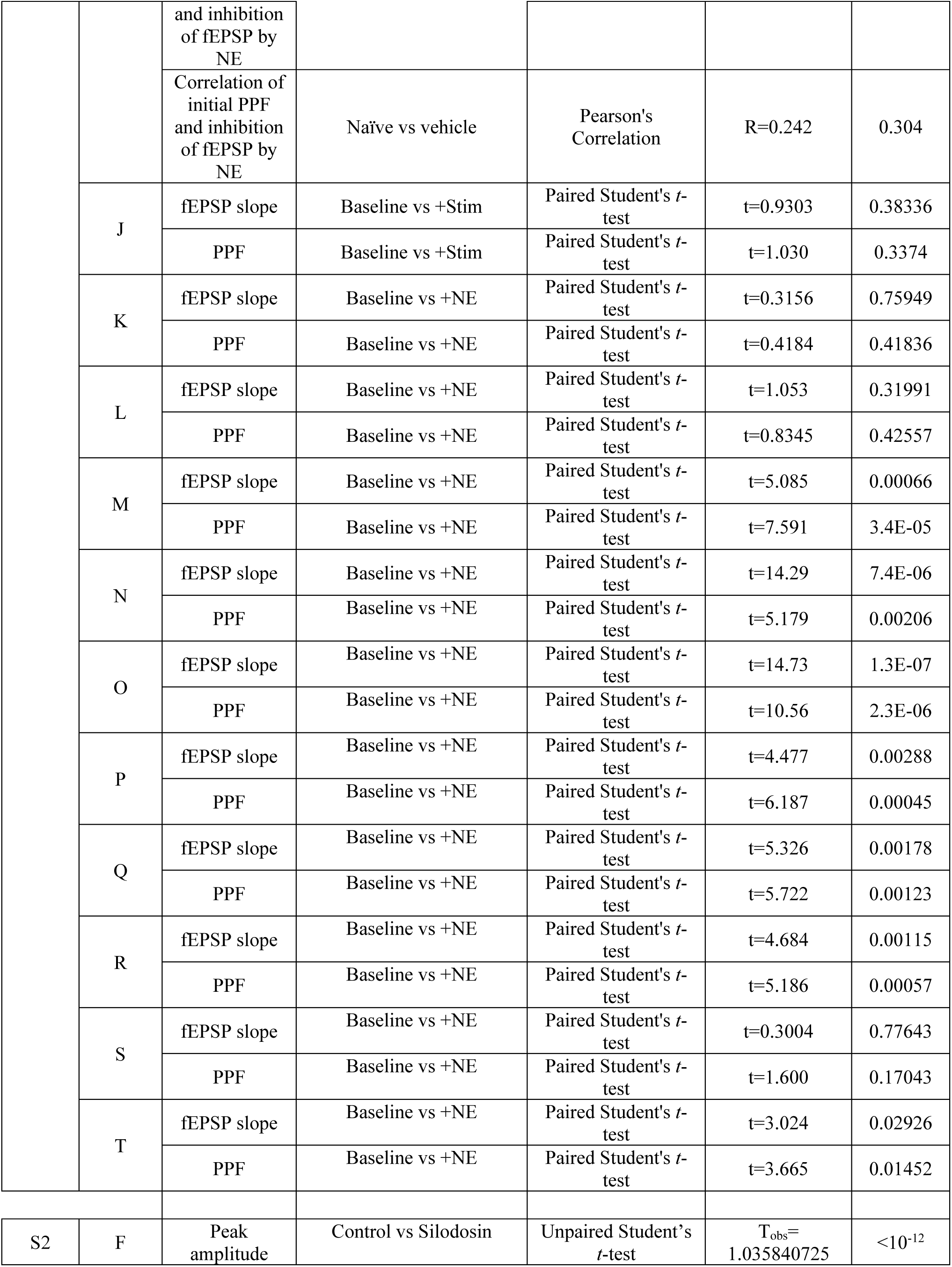
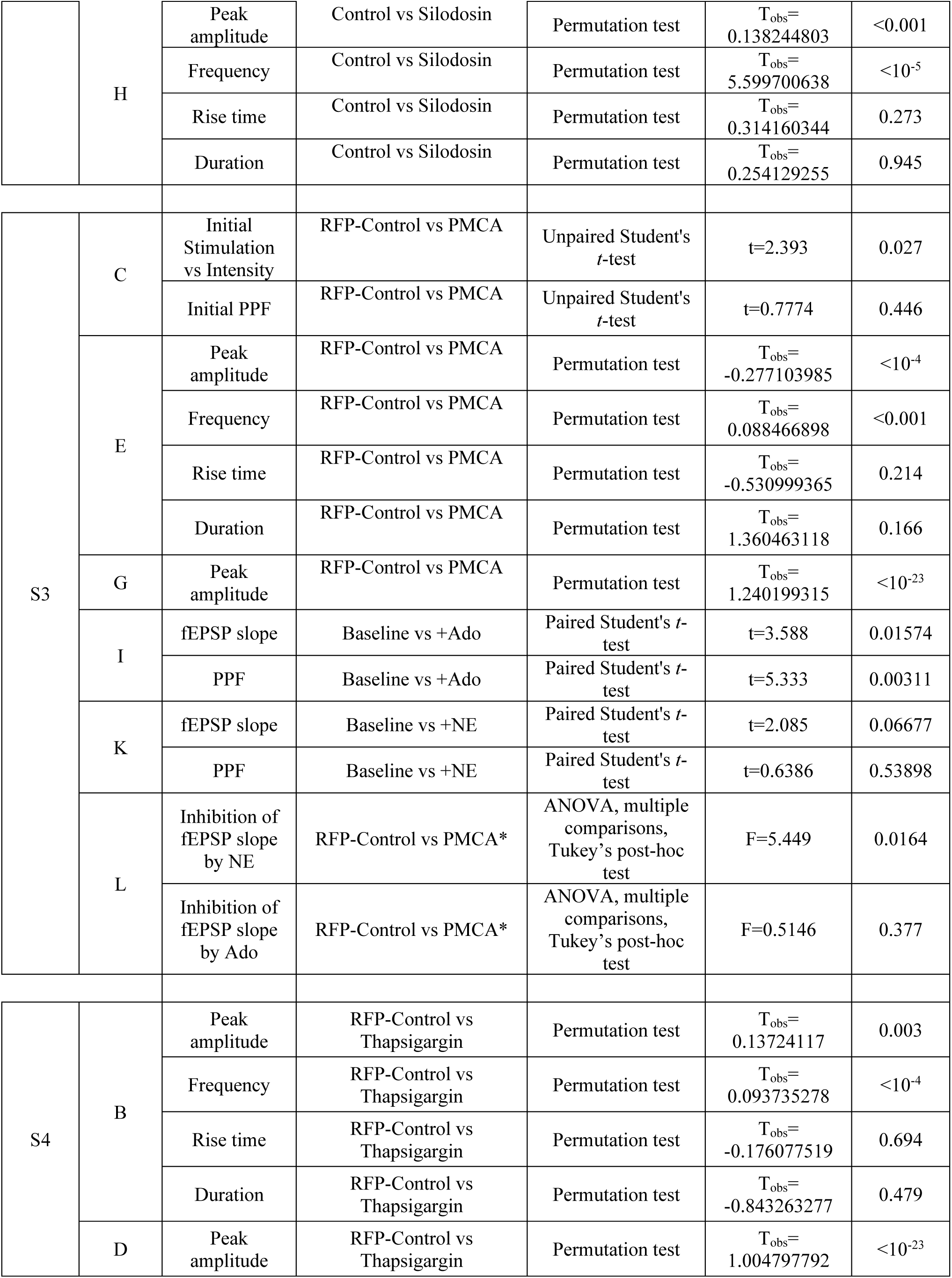
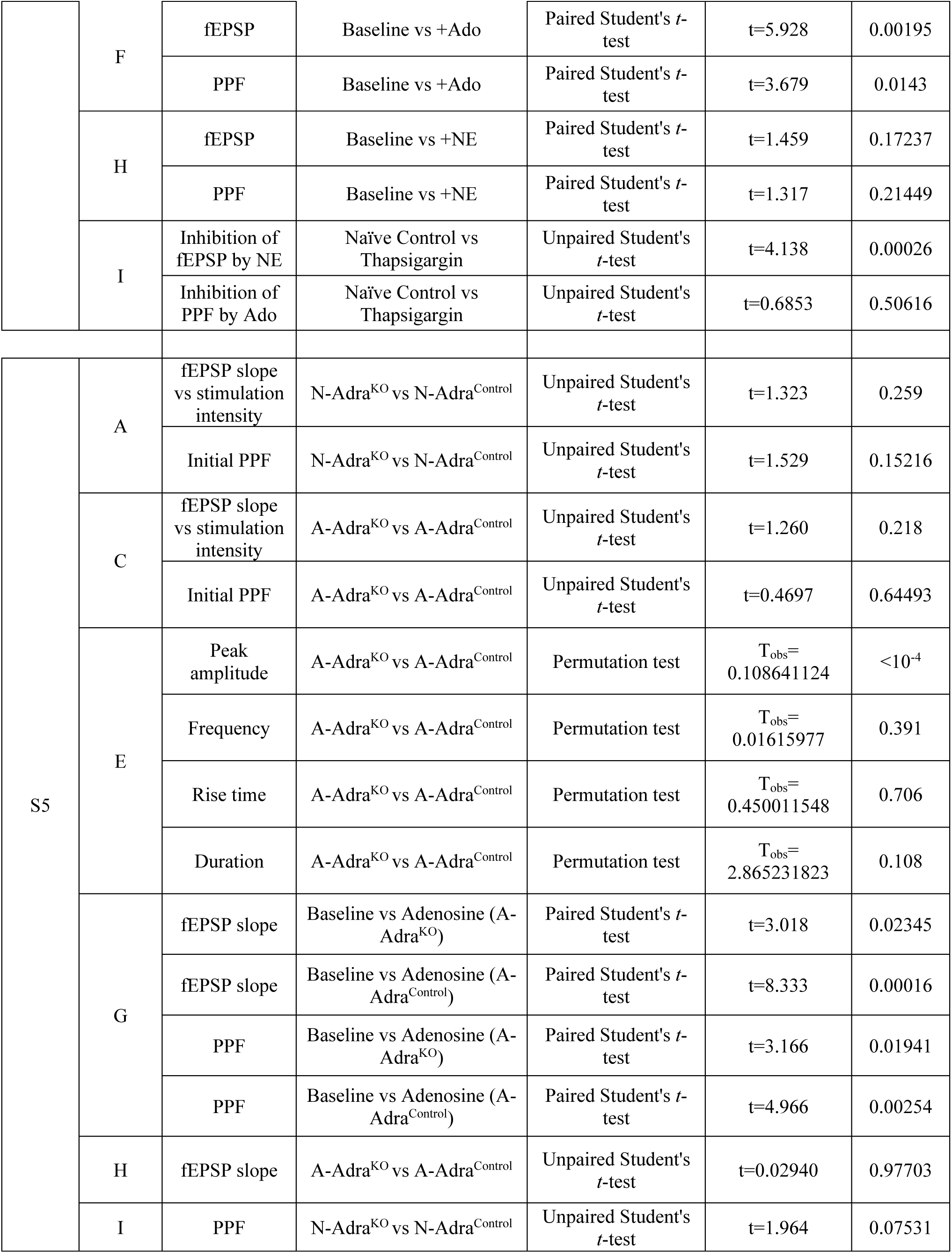
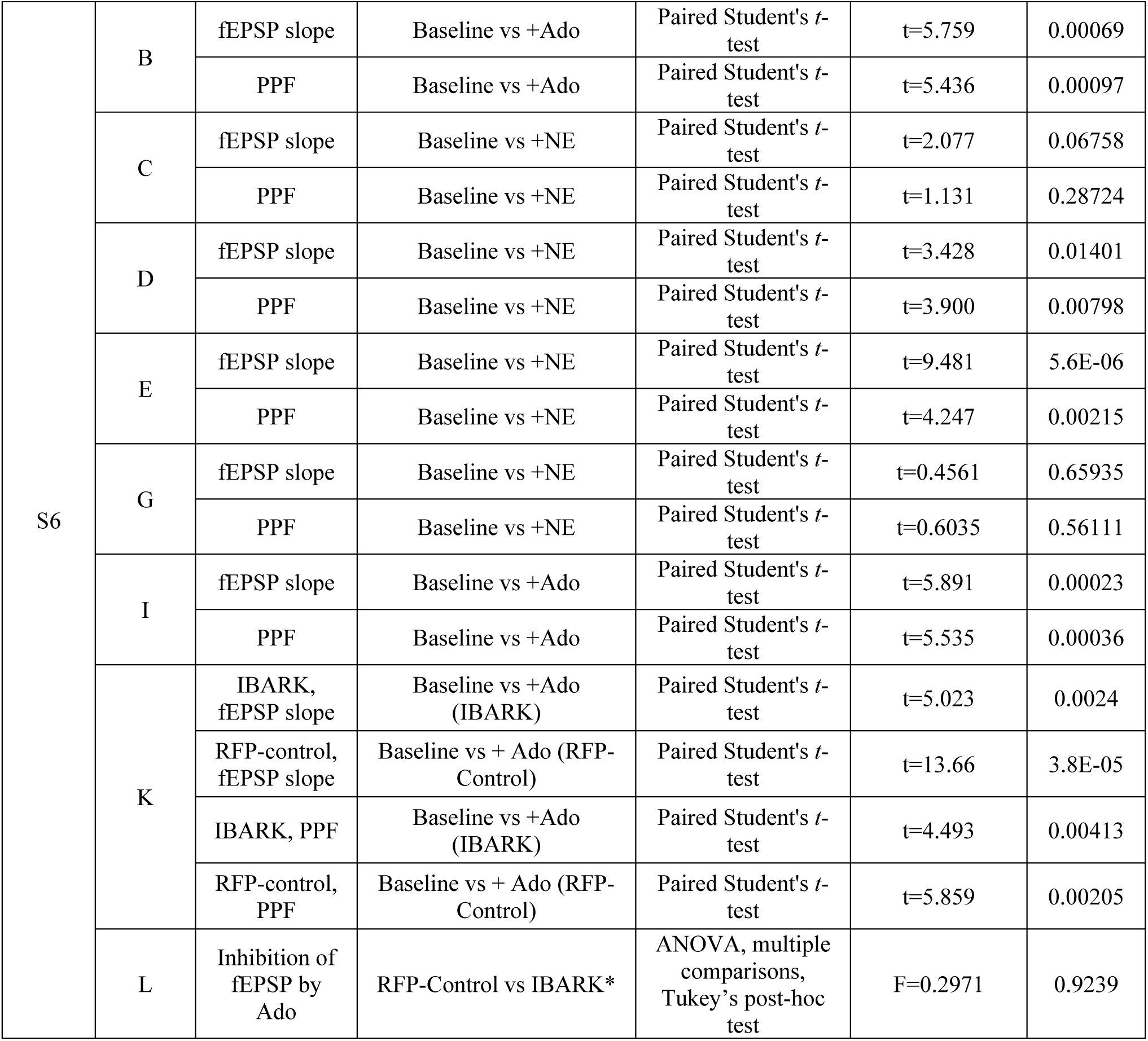
Summary of statistical tests, p values, and statistical test values for all experiments illustrated in figures. * indicates that experiments were compared across RFP-Control, iβARK, and PMCA conditions.

**Table S2:**
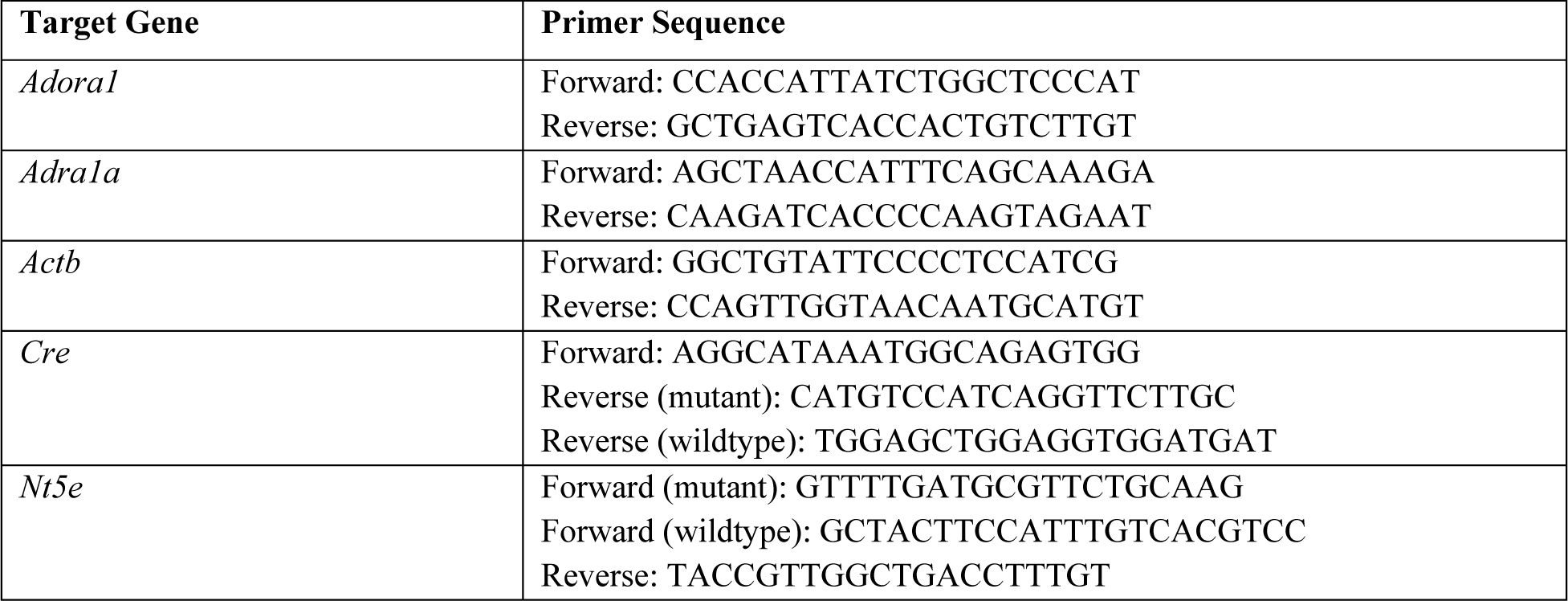
Primer sequences used for genotyping and validation of recombination. All primers listed in 5’ to 3’.

**Table S3:** Primary antibodies used for immunofluorescence experiments.

| Target | Host | Vendor | Product Number |
| --- | --- | --- | --- |
| GFAP | Chicken | Abcam | AB4674 |
| NeuN | Guinea Pig | Millipore Sigma | ABN90 |
| Tyrosine Hydroxylase | Rabbit | ThermoFisher | PA5-85167 |
| GFP | Rabbit | Abcam | AB290 |
| RFP | Rabbit | Abcam | AB62341 |

**Table S4:** Secondary antibodies used for immunofluorescence experiments.

| Host | Alexa Fluorophore | Vendor | Product Number |
| --- | --- | --- | --- |
| Goat-anti-Rabbit | 488 | ThermoFisher | A-11034 |
| Goat-anti-Rabbit | 568 | ThermoFisher | A-11011 |
| Goat-anti-Chicken | 488 | ThermoFisher | A-11039 |
| Goat-anti-Chicken | 647 | ThermoFisher | A-32933 |
| Goat-anti-Guinea Pig | 647 | ThermoFisher | A-21450 |

